# A neural basis of probabilistic computation in visual cortex

**DOI:** 10.1101/365973

**Authors:** Edgar Y. Walker, R. James Cotton, Wei Ji Ma, Andreas S. Tolias

## Abstract

Bayesian models of behavior suggest that organisms represent uncertainty associated with sensory variables. However, the neural code of uncertainty remains elusive. A central hypothesis is that uncertainty is encoded in the population activity of cortical neurons in the form of likelihood functions. We studied the neural code of uncertainty by simultaneously recording population activity from the primate visual cortex during a visual categorization task in which trial-to-trial uncertainty about stimulus orientation was relevant for the decision. We decoded the likelihood function from the trial-to-trial population activity and found that it predicted decisions better than a point estimate of orientation. This remained true when we conditioned on the true orientation, suggesting that internal fluctuations in neural activity drive behaviorally meaningful variations in the likelihood function. Our results establish the role of population-encoded likelihood functions in mediating behavior, and provide a neural underpinning for Bayesian models of perception.

---

When making perceptual decisions, organisms often benefit from representing uncertainty about sensory variables. More specifically, the theory that the brain performs Bayesian inference— which has roots in the work of Laplace^1^ and von Helmholtz^2^—has been widely used to explain human and animal perception^3–6^. At its core lies the assumption that the brain maintains a statistical model of the world and when confronted with incomplete and imperfect information, makes inferences by computing probability distributions over task-relevant world state variables (e.g. direction of motion of a stimulus). In spite of the prevalence of Bayesian theories in neuroscience, evidence to support them stems primarily from behavioral studies (e.g.^7,8^). Consequently, the manner in which probability distributions are encoded in the brain remains unclear, and, thus, the neural code of uncertainty is unknown.

It has been hypothesized that a critical feature of the neural code of uncertainty, which is shared throughout the sensory processing chain in the neocortex, is that the same neurons that encode a specific world state variable (e.g. stimulus orientation in V1) also encode the uncertainty about that variable (Fig. 1a). Therefore neurons multiplex both a point estimate of a sensory variable and the associated uncertainty about it^9,10^. Specifically, according to the probabilistic population coding (PPC) hypothesis^9,10^, inference in the brain is performed by inverting a generative model of neural population activity. Under this coding scheme, neural populations in V1, for example, that encode stimulus orientation also encode the associated uncertainty in the form of the sensory likelihood function—the probability of observing a given pattern of neural activity across hypothesized stimulus values^9,11,12^. The form of the likelihood function is related to the probability distribution describing neural variability (“noise”) for a given stimulus. A sensory likelihood function is often unimodal^13,14^, and its width could in principle serve as a measure of the sensory uncertainty about the stimulus. Whether the brain uses this particular uncertainty quantity in its decisions is unknown. Alternatively, it may be the case that the neural population that encodes an estimate of a sensory variable (e.g. stimulus orientation in V1) does not carry information about the associated uncertainty (Fig. 1b).

**Figure 1:**
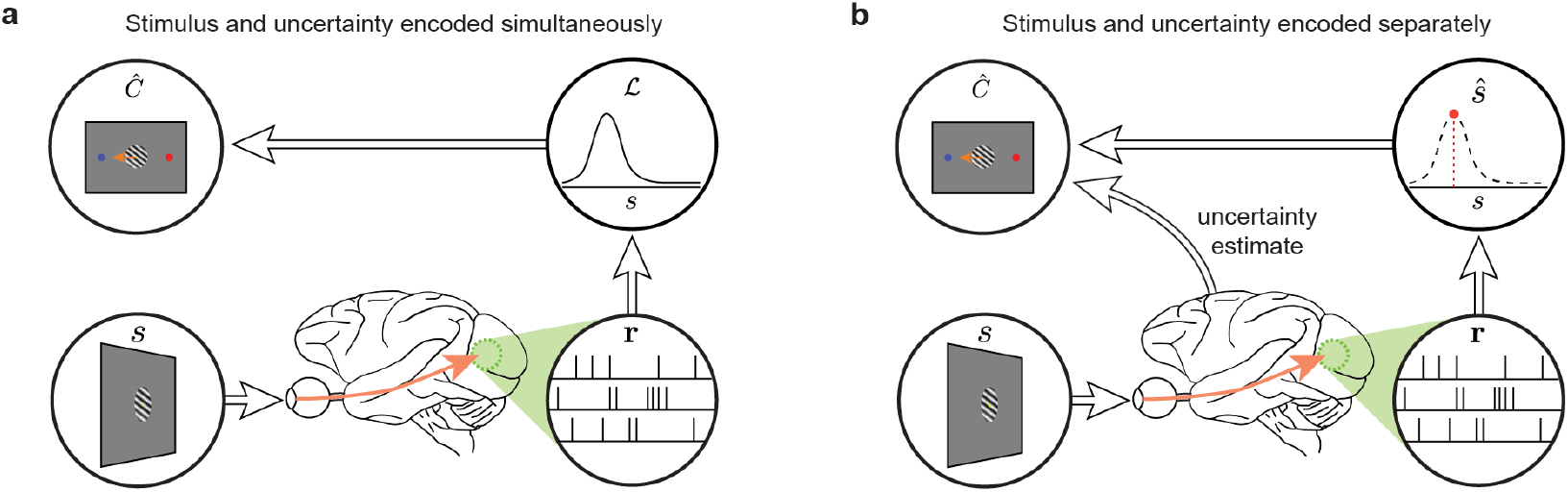
Alternative models of uncertainty information encoding. **a**, The recorded cortical population **r** responding to sensory stimulus *s* encodes stimulus estimate and uncertainty simultaneously in the form of likelihood function 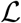 which is subsequently used in making a decision *Ĉ* as the subject performs a visual classification task. **b**, The recorded cortical population only encodes a point estimate of the stimulus *ŝ* while an estimate of the sensory uncertainty is made by other (unrecorded) cortical populations. The information is subsequently combined to lead to the subject’s decision *Ĉ*.

We recorded the activity of V1 cortical populations while monkeys performed a visual classification task in which the trial-by-trial uncertainty information is beneficial to the animal^15^. To decode the trial-by-trial likelihood functions from the V1 population responses, we developed a novel technique based on deep learning^16,17^. Importantly, we performed all analyses conditioned on the contrast—an overt driver of uncertainty—and performed further orientation-conditioned analyses to isolate the effect of random fluctuations in the decoded likelihood function on behavior. We found that using the trial-to-trial changes in the shape of the likelihood function allowed us to better predict the behavior than using a likelihood function with a fixed shape shifted by a point estimate. Therefore, we provide the first evidence that in perceptual decision-making, the same cortical population that encodes a sensory variable also encodes its trial-by-trial sensory uncertainty information, which is used to mediate perceptual decisions, consistent with the theory of PPC.

## Results

### Behavior

Two Rhesus macaques *(Macacca mulatta*) were trained on an orientation classification task designed such that the optimal performance required the use of trial-by-trial uncertainty. On each trial, one of two stimulus classes (*C* = 1 or *C* = 2) was chosen at random with equal probability. Each class was defined by a Gaussian probability distribution over the orientation. The two distributions shared the same mean but had different standard deviations (Fig. 2a). An orientation was drawn from the distribution belonging to the selected class, and a drifting grating stimulus with that orientation was then presented to the animal (Fig. 2b). In a given recording session, at least three distinct contrasts were selected at the beginning of the session, and on each trial, one of these values was randomly selected.

**Figure 2:**
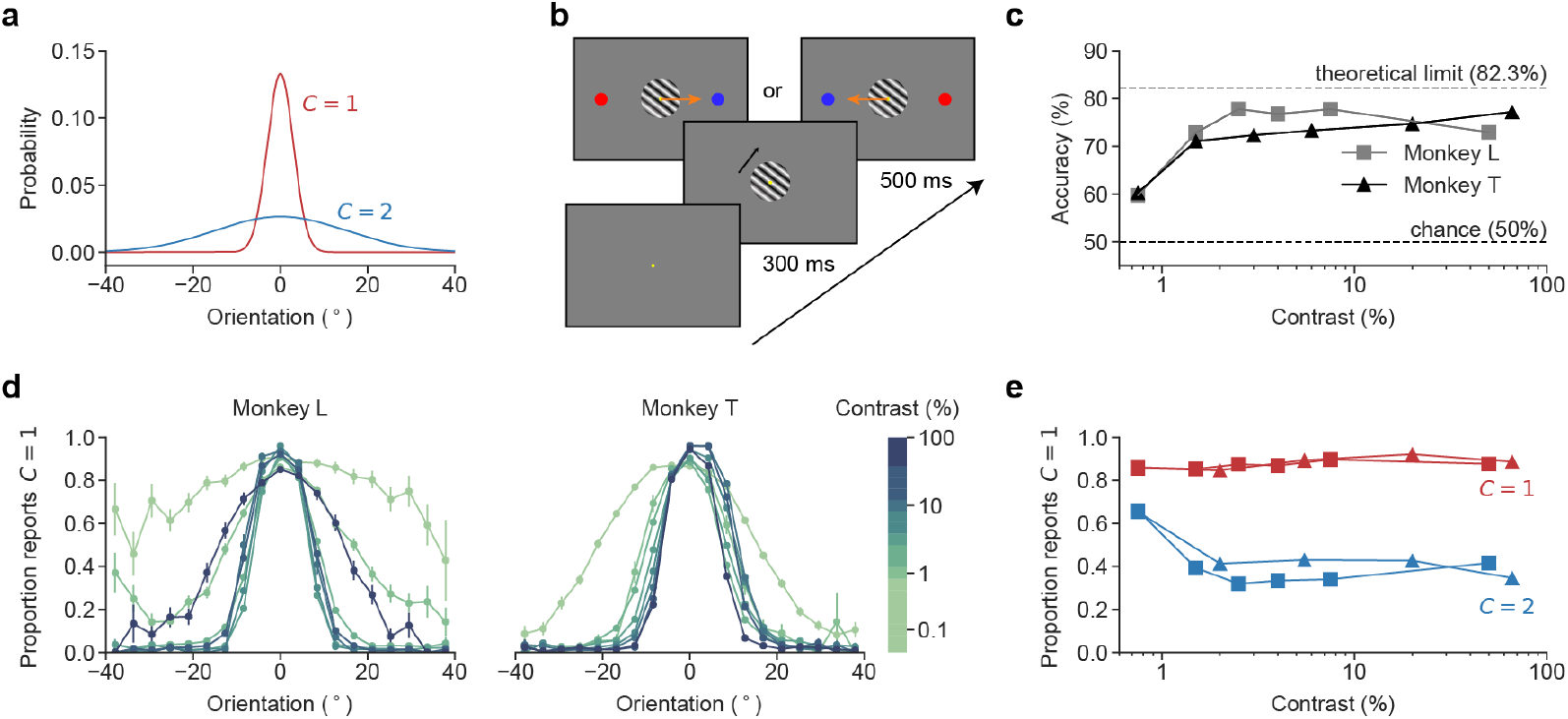
Behavioral task. **a**, The stimulus orientation distributions for the two classes. The two distributions shared the same mean (*μ* = 0°) but differed in their standard deviations (*σ*_1_ = 3° and *σ*_2_ = 15°). **b**, Time course of a single trial. The subject fixated onto the fixation target for 300 ms before a drifting grating stimulus was shown. After 500 ms of stimulus presentation, the subject broke fixation and saccaded to one of the two colored targets to indicate their class decision (color matches class color in **a**). The left-right configuration of the colored targets was chosen at random for each trial. **c**, Performance of the two monkeys on the task across stimulus contrast. “Theoretical limit” corresponds to the performance of an ideal observer with no observation noise. **d**, Psychometric curves. Each curve shows the proportion of trials on which the monkey reported *C* = 1 as a function of stimulus orientation, computed from all trials within a single contrast bin. All data points are means and error bars indicate standard error of the means. e, Class-conditioned responses. For each subject, the proportions of *C* =1 reports is shown across contrasts, conditioned on the ground-truth class: *C* =1 (red) and *C* = 2 (blue). The symbols have the same meaning as in **c**.

In our previous study^15^, we designed this task so that an optimal Bayesian observer would incorporate the trial-by-trial sensory uncertainty about stimulus orientation in making classification decisions. Indeed, decisions of both humans and monkeys seemed to utilize trial-by-trial uncertainty about the stimulus orientation. In the current study, one of the two monkeys (Monkey L) was the same monkey that participated in the previous study and thus has been shown to have learned the task well. A second monkey (Monkey T) was also trained on the task and closely matched the performance of Monkey L (Fig. 2c). Both animals had psychometric curves displaying the expected strong dependence on both contrast and orientation (Fig. 2d,e).

In our analyses, we grouped the trials with the same contrast within the same session and refer to such a group as a “contrast-session”.

### Decoding likelihood function from V1

Each monkey was implanted with a chronic multi-electrode (Utah) array in the parafoveal primary visual cortex (V1) to record the simultaneous cortical population activity as the subjects performed the orientation classification task (Fig. 3a).

**Figure 3:**
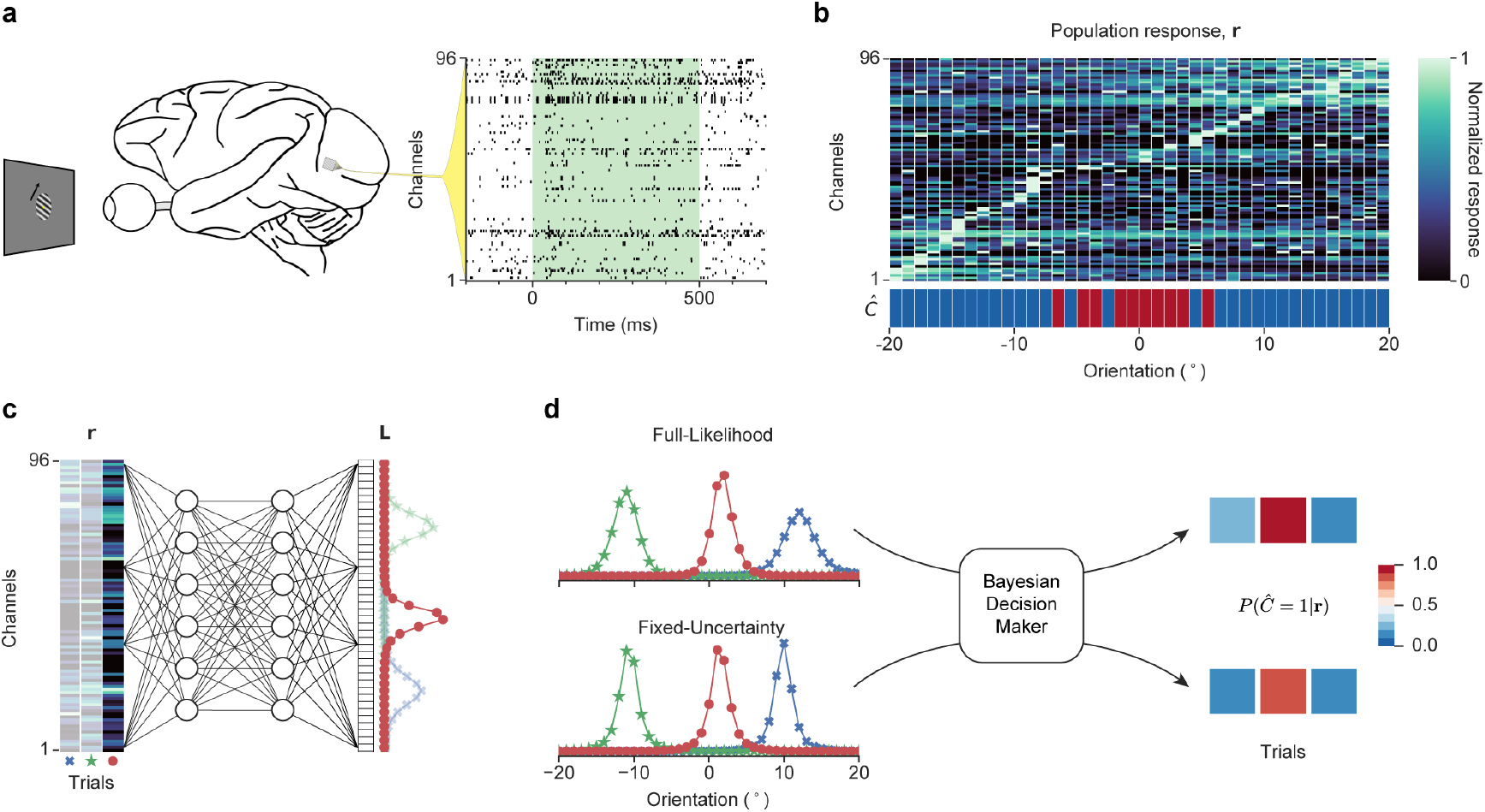
Encoding and decoding of the stimulus orientation. **a**, An example of 96 channels spike traces from a single trial (Monkey T). The vector of spike counts, **r**, was accumulated over the pre-saccade stimulus presentation period (time 0-500 ms, green shade). **b**, The population response for the selected trials from a single contrast-session (Monkey T, 64% contrast). Column: a population response **r** on a trial randomly drawn from the trials falling into a specific orientation bin. Row: a response from a single channel. For visibility, the channel’s responses are normalized to the maximum response across all trials. The channels were sorted by the preferred orientation of the channel. Subject’s class decision is indicated by red and blue color patches for *Ĉ* = 1 and *Ĉ* = 2, respectively. **c**, A schematic of a DNN for the Full-Likelihood decoder, mapping **r** to the decoded likelihood function **L**. All likelihood functions are area-normalized. **d**, Two models of likelihood decoder *M*. In the *Full-Likelihood decoder*, the likelihood L was decoded without any constraints on the shape. In the *Fixed-Uncertainty decoder*, all decoded likelihood functions shared the same shape but differed in the location of the center based on the population response. For both decoders, the resulting likelihood functions were fed into parameterized Bayesian decision models to yield the decision prediction *p*(*Ĉ* = 1 |**r**, *M*).

A total of 61 and 71 sessions were analyzed from Monkey L and Monkey T for a total of 110,695 and 192,631 trials, respectively (Supplementary Fig. 1). In each recording session, up to 96 channels were recorded. On each trial and for each channel, we computed the total number of spikes that occurred during the 500 ms of stimulus presentation preceding the decision-making cue (Fig. 3a), yielding a vector of population responses **r** used in the subsequent analyses (Fig. 3b).

Existing computational methods for decoding the trial-by-trial likelihood function from the cortical population activities typically make strong parametric assumptions about the stimulus conditioned distribution of the population response (i.e. the generative model of the population response). For example, population responses to a stimulus can be modeled as an independent Poisson distribution, allowing each recorded unit to be characterized by a simple tuning curve (which may be further parameterized)^14,18–22^. While this simplifying assumption makes computing the trial-by-trial likelihood function straightforward, disregarding potential correlations among the units in population responses (i.e. noise correlations and internal brain state fluctuations^23–28^) can lead to biased estimates of the likelihood function and limits the generality of this approach. While more generic parametric models—such as Poisson-like distributions—of population distributions have been proposed^9,10,15,29,30^, they still impose restrictive assumptions.

We devised a technique based on deep learning to decode the trial-by-trial likelihood function from the V1 population response. This neural network-based likelihood decoder allows us to approximate the information that can be extracted about the stimulus orientation from the cortical population responses. The network was not used as a model of how the rest of the brain extracts and processes the information present in the population, but rather to decode it and demonstrate that it is used behaviorally.

We trained a fully connected deep neural network (DNN)^17^ to predict the per-trial likelihood function 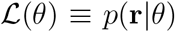 over stimulus orientation *θ* from the vectorized population response **r** (Fig. 3c; for details on the network architecture, training objective, and hyperparameter selection see Methods and Supplementary Table 1). A separate network was trained for each contrast-session and no behavioral data were utilized in training the DNN.

Using a DNN to decode the likelihood function avoids the restrictive parametric assumptions described above and provides a strictly more flexible method, often capturing decoding under known distributions as a special case (Supplementary Fig. 2). We demonstrate this by showing the DNN can recover the ground-truth likelihood function from simulated responses sampled from known distributions (Supplementary Fig. 3; refer to Methods for the simulation details).

The likelihood functions decoded by the DNNs exhibited the expected dependencies on the overt drivers of uncertainty such as contrast (Fig. 4a-c): the width of the likelihood function is higher at lower contrast (Fig. 4d).

**Figure 4:**
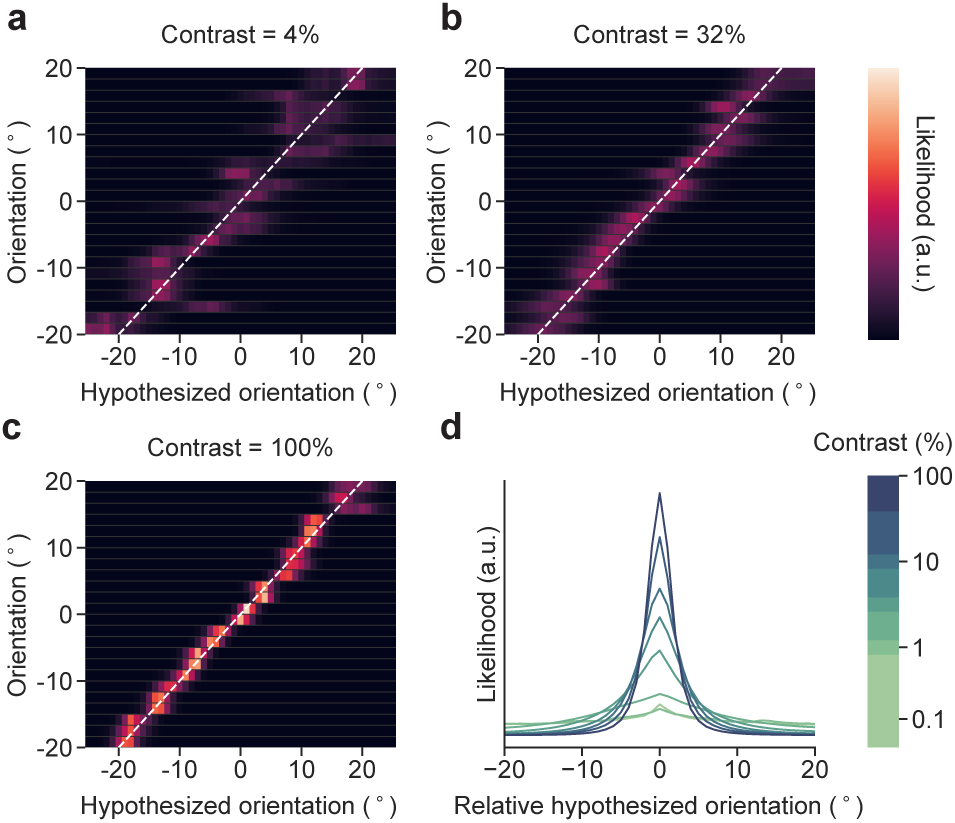
Likelihood functions decoded by the trained neural networks. **a-c**, Example decoded likelihood functions from three contrast-sessions from Monkey T. Each row represents the decoded likelihood function over the hypothesized orientation for a randomly selected trial within the specific orientation bin. All likelihood functions are area-normalized. Brighter colors correspond to higher values of the likelihood function. **d**, Average likelihood function by contrast. On each trial, the likelihood function was shifted such that the mean orientation of the normalized likelihood function occurred at 0°. The centered likelihood functions were then averaged across all trials within the same contrast bin.

### Trial to trial uncertainty improves behavioral predictions

To assess whether the uncertainty decoded from population responses in the form of sensory likelihood functions mediate the behavioral outcome (perceptual decision) as we hypothesized, it is critical that we appropriately condition the analysis on the stimulus. To illustrate the impor-tance of conditioning on the stimulus to determine if the decoded likelihood function mediates perceptual decisions, consider a typical perceptual decision-making task (like ours) (Supplementary Fig. 4) where the subject views a stimulus s, which elicits a population response r, for example in V1. Here, by “stimulus”, we refer collectively to all aspects of a visual stimulus, such as its contrast and orientation. Stimulus information is eventually relayed to decisionmaking areas (e.g. prefrontal cortex), leading the animal to make a classification decision *Ĉ*. We decode the likelihood function 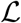 from the recorded population activity **r**. Because variation in the stimulus (e.g. orientation or contrast) across trials can drive variation both in the decoded likelihood function and in the animal’s decision, one may find a significant relationship between 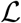 and *Ĉ*, even if the likelihood function estimated from the recorded population **r** does not mediate the decision. When the stimulus is fixed, random fluctuations in the population response **r** can still result in variations in 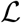. If the likelihood function truly mediates the decision, we expect that such variation in 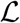 would account for variation in *C*. Therefore, to demonstrate that the likelihood 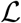 mediates the decision *C*, it is imperative to show a correlation between 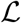 and *Ĉ* conditioned on the stimulus *s*.

As we varied the stimulus contrast from trial to trial in our task, the expected uncertainty about the stimulus orientation varied, and one would expect the monkeys to represent and make use of their trial-by-trial sensory uncertainty in making decisions. However, we make a much stronger claim here: even at a fixed contrast, because of random fluctuations in the population response^31,32^, we predict (1) the uncertainty encoded in the population, that is, the likelihood function, will still fluctuate from trial to trial, and (2) the effect of such fluctuations will manifest in the monkey’s decisions on a trial-by-trial basis.

We tested this prediction by fitting, separately for each contrast-session, the following two decision models and comparing their performance in predicting the monkey’s trial-by-trial decisions: (1) a Full-Likelihood Model, which utilizes the trial-by-trial uncertainty information decoded from the population response in the form of the likelihood function obtained from the neural-network based likelihood decoder (Full-Likelihood decoder) described above (Fig. 3d), and (2) a Fixed-Uncertainty Model, which utilizes an alternative neural-network based likelihood decoder (Fixed-Uncertainty decoder) that learns a single, fixed-shape likelihood function whose location is shifted from trial to trial based on the population response (Supplementary Fig. 5). The Fixed-Uncertainty Model captures the alternative hypothesis in which the recorded sensory population only encodes a point estimate of the sensory variable (i.e. mean of the likelihood function) and the estimate of the sensory uncertainty is encoded elsewhere, signified by the fixed shape of the likelihood function fitted for each contrast-session under this model (Fig. 1b). Generally, the likelihood function decoded by Fixed-Uncertainty decoder closely approximated the likelihood function decoded by the Full-Likelihood decoder (Supplementary Fig. 5). We use the term decoder for the DNN that returns estimated likelihood functions, and the term decision model for the mapping from likelihood function to decision.)

In both models, the decoded likelihood functions were fed into the Bayesian decision maker to yield trial-by-trial predictions of the subject’s decision in the form of *p*(*Ĉ*|**r**, *M*), or the likelihood of subject’s decisions *Ĉ* conditioned on the population response **r** and the decision model *M*. The Bayesian decision maker computed the posterior probability of each class and used these to produce a stochastic decision. The means of the class distributions assumed by the observer, the class priors, the lapse rate, and a parameter to adjust the exact decision-making strategy were used as free parameters (Supplementary Fig. 6, refer to Methods for details). The model parameters were fitted by maximizing the total log likelihood over all trials for each contrast-session 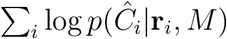. The fitness of the models was assessed through crossvalidation, and we reported mean and total log likelihood of the models across all trials in the test set.

Both models incorporated trial-by-trial changes in the point estimate of the stimulus orientation (e.g. the mean of the likelihood function) and only differed in whether they contained additional uncertainty information about the stimulus orientation carried by the trial-by-trial fluctuations in the shape of the likelihood function decoded from the same population that encoded the point estimate. We use the term “shape” to refer to all aspects of the likelihood function besides its mean, including its width. If the fluctuations in the shape of the likelihood function truly captured the fluctuations in the sensory uncertainty as represented and utilized by the animal, one would expect the Full-Likelihood Model to yield better trial-by-trial predictions of the monkey’s decisions than the Fixed-Uncertainty Model.

We observed that both models predicted the monkey’s behavior well across all contrasts (Supplementary Fig. 7), reaching up to 90% accuracy. We also observed that the performance of the decision models using likelihood functions that were decoded by the neural networks was superior to the models using likelihood functions that were decoded with more traditional parametric generative models (independent Poisson distribution and Poisson-like distribution) (Supplementary Fig. 8; refer to Methods for details). The Full-Likelihood Model consistently outperformed the Fixed-Uncertainty Model across contrasts and for both monkeys (Fig. 5a,b; trial log likelihood differences between the Full-Likelihood and Fixed-Uncertainty Model: Monkey L: paired t-test, *t*(110694) = 11.06, *p* < 0.001, *δ*_total_ = 11.0 × 10^2^ and Monkey T: *t*(192610) = 11.03, *p* < 0.001, *δ*_total_ = 11.3 × 10^2^; *δ*_total_ is the total log likelihood difference across all trials). This result shows that the trial-by-trial fluctuations in the shape of the likelihood function are informative about the monkey’s trial-by-trial decisions, demonstrating that decision-relevant sensory uncertainty information is contained in population responses that can be captured by the shape of the full likelihood function. This finding in turn strongly supports the hypothesis that visual cortex encodes stimulus uncertainty through the shape of the full likelihood function on a trial-by-trial basis.

**Figure 5:**
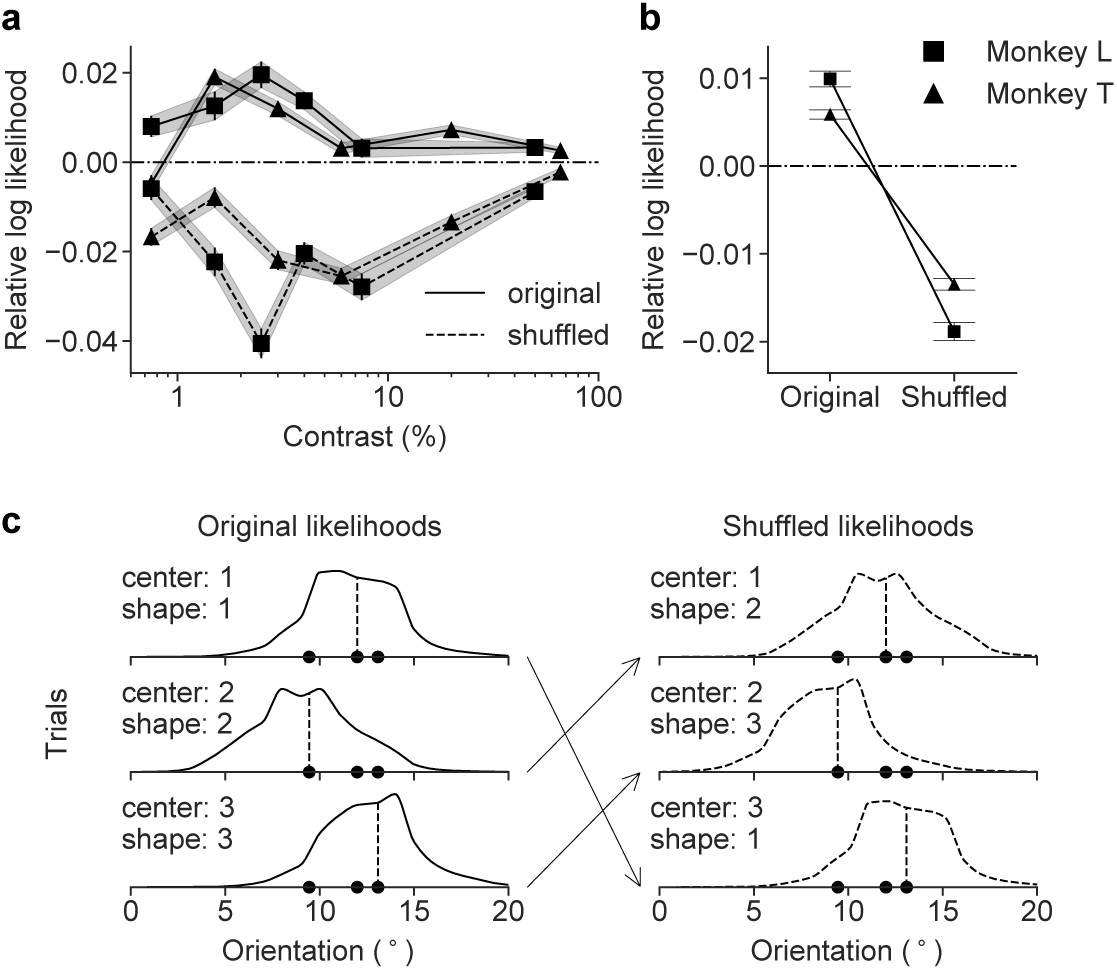
Model performance. **a**, Average trial-by-trial performance of the Full-Likelihood Model relative to the Fixed-Uncertainty Model across contrasts, measured as the average trial difference in the log likelihood. The results for the original (unshuffled) and the shuffled data are shown in solid and dashed lines, respectively. The squares and triangles mark Monkey L and T, respectively. **b**, Relative model performance summarized across all contrasts. Performance on the original and the shuffled data is shown individually for both monkeys. The difference between the Full-Likelihood and Fixed-Uncertainty Models was significant with *p* < 0.001 for both monkeys, and on both the original and the shuffled data. Furthermore, the difference between the Full-Likelihood Model on the original and the shuffled data was significant (*p* < 0.001 for both monkeys). For **a** and **b**, all data points are means, and error bar/shaded area indicate standard error of the means. **c**, Shuffling scheme for three example trials drawn from the same stimulus orientation bin. Shuffling maintains the means but swaps the shapes of the likelihood functions.

We repeated this analysis after splitting the data into the first and second 250ms of stimulus presentation. We found a similar improvement for the Full-Likelihood model over the Fixed-Uncertainty model in both periods (Supplementary Fig. 9).

We next asked how meaningful our effect sizes (model performance differences) are. To answer this question, we simulated the monkey’s responses across all trials and contrast-sessions taking the trained Full-Likelihood Model to be the ground truth, and then retrained the Bayesian decision makers in the Full-Likelihood Model and the Fixed-Uncertainty Model from scratch on the simulated data. This approach yields a theoretical upper bound on the observable difference between the two models if the Full-Likelihood Model was the true model of the monkeys’ decision-making process.

We observed that the expected total upper bound log likelihood differences between the Full-Likelihood Model and the Fixed-Uncertainty Model of (37.1 ± 1.5) × 10^2^ and (36.0 ± 1.3) × 10^2^ based on the simulations (representing mean ± standard deviation across 5 repetitions of simulation for Monkey L and T, respectively) were larger but in the same order of the magnitude as the observed model performance differences (11.0 × 10^2^ and 11.3 × 10^2^ total log likelihood differences across all trials for Monkey L and T, respectively), suggesting that our effect sizes are meaningful and that the Full-Likelihood Model is a reasonable approximate description of the monkey’s true decision-making process (Supplementary Fig. 10).

### Stimulus dependent changes in uncertainty

We observed that for some contrast-sessions, the average width of the likelihood function showed a dependence on the stimulus orientation (Supplementary Figure. 11). By design, the Fixed-Uncertainty Model cannot capture this stimulus dependent change in uncertainty, which could contribute to it under-performing the Full-Likelihood Model (Supplementary Fig. 4).

To rule this out, we shuffled the shapes of the decoded likelihood functions across trials within the same orientation bin, separately for each contrast-session. This shuffling preserves the average stimulus-dependent change in uncertainty and trial-by-trial correlation between the mean of the likelihood function and the decision (Fig. 5c), while removing the trial-by-trial correlation between the shape of the likelihood function and the behavioral decision conditioned on the stimulus orientation.

By design, the Fixed-Uncertainty Model makes identical predictions on the original and the shuffled data. If the Full-Likelihood Model outperformed the Fixed-Uncertainty Model simply because it captured spurious correlations between the stimulus orientation and the shape of the likelihood function, then it should outperform the Fixed-Uncertainty model by the same amount on the shuffled data. However, if the better behavioral predictions come from the trial-by-trial fluctuations in the likelihood shape as we hypothesized, one would expect this difference to disappear on the shuffled data. Indeed, the shuffling of the likelihood function shapes abolished the improvement in prediction performance that the Full-Likelihood Model had over the Fixed-Uncertainty Model. In fact, the Full-Likelihood Model consistently underperformed the Fixed-Uncertainty Model on the shuffled data (Fig. 5a,b; trial log likelihood difference between the Full-Likelihood Model and the Fixed-Uncertainty Model on the shuffled data: Monkey L: paired t-test *t*(110694) = −18.44, *p* < 0.001, *δ*_total_ = −20.9 × 10^2^ and Monkey T: *t*(192610) = −20.15, *p* < 0.001, *δ*_total_ = −25.9 × 10^2^; *δ*_total_ is the total log likelihood difference across all trials). Therefore, there were significant performance differences in Full-Likelihood Model between the unshuffled and shuffled data (trial log likelihood difference: Monkey L: paired t-test *t*(110694) = 33.34, *p* < 0.001, *δ*_total_ = 31.9 × 10^2^ and Monkey T: *t*(192610) = 34.52, *p* < 0.001, *δ*_total_ = 37.2 × 10^2^).

To confirm our effect sizes were appropriate, we again compared these values to those obtained from simulations in which we took the Full-Likelihood Model to be the ground truth (Supplementary Fig. 10). The simulations yielded total log likelihood differences of the Full-Likelihood Model between the unshuffled and shuffled data of (36.2 ± 2.2) × 10^2^ (Monkey L) and (40.7 ± 1.5) × 10^2^ (Monkey T) (mean ± standard deviation across 5 repetitions), similar in magnitude to the observed values.

Taken together, the shuffling analyses show that the better prediction performance of the Full-Likelihood Model is not due to the confound between the stimulus and the likelihood shape. We conclude that the trial-by-trial likelihood function decoded from the population represents behaviorally relevant stimulus uncertainty information, even when conditioned on the stimulus.

### Attribution analysis

To assess whether the same population encoding the best point estimate (i.e. mean of the likelihood function) also encoded the uncertainty regarding that estimate (i.e. shape of the likelihood function), as we hypothesized to be the case, we performed attribution analysis^33^ on the trained Full-Likelihood decoder. Through this analysis, we ask how much of the changes in either (1) the mean of the likelihood *μ_L_* (i.e. surrogate for the best point estimate) or (2) the standard deviation of the likelihood function *σ_L_* (i.e. surrogate measure of the uncertainty) can be attributed back to each input multiunit, yielding attribution **A**_*μ*_ and **A**_*σ*_, respectively. The question of feature attribution is a very active field of research in machine learning, and multiple methods of attribution computation exist^33–35^. Here we have selected three different methods of computing attribution scores: saliency maps^34^, gradient × input^33^, and DeepLIFT^35^ (refer to Methods for the details of attribution computation).

We observed that across all three attribution methods, multiunits with high *μ_L_* attribution tended to have high *σ_L_* attribution, and vice versa, giving rise to high degree of correlation between *A_μ_* and *A_σ_* (Fig. 6a). If distinct subpopulations were involved in encoding the point estimate and the uncertainty as found in the likelihood function, we would have expected multiunits with a high *μ_L_* attribution score to have a low *σ_L_* attribution score, and vice versa, there-fore leading to negative correlation between *A_μ_* and *A_σ_*. However, we observed that across all contrast-sessions from both monkeys, *A_μ_* was strongly positively correlated with *A_σ_* regardless of the exact attribution method used, suggesting that the highly overlapping subpopulations are involved in encoding both the point estimate and the uncertainty of the likelihood function, as we hypothesized would be the case (Fig. 6b-d).

**Figure 6:**
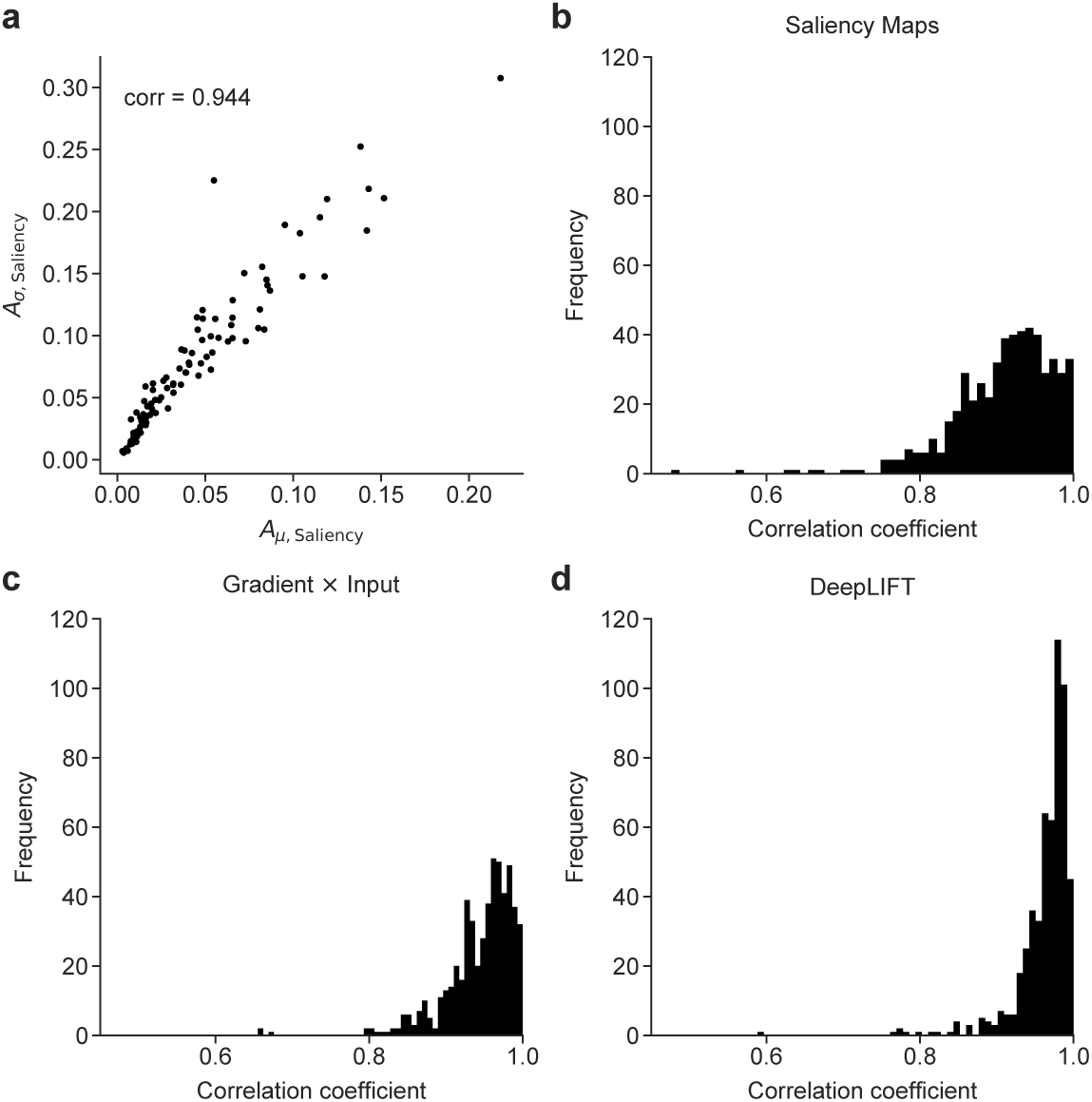
Attribution analysis for means and standard deviations of the likelihood functions. **a**, Attribution of 96 input multiunits to the likelihood mean *A_μ,Saliency_* vs. standard deviation *A_σ,Saliency_* computed based on saliency maps for an example contrast session (Monkey T, 32% contrast). b-d, Distribution of correlation coefficients between *A_μ_* and *A_σ_* for multi units across all contrast sessions for both monkeys, computed based on different attribution methods.

## Discussion

Given the stochastic nature of the brain, repeated presentations of identical stimuli elicit variable responses. The covariation between neuronal activity fluctuations and perceptual choice has been studied extensively at the level of single neurons, originating with the pioneering work of Campbell & Kulikowski^36^ and Britten et al.^37^. Here, we go beyond this literature by examining the hypothesis that the brain takes into account knowledge of the form of neural variability in order to build a belief over the stimulus of interest on each trial. This belief is captured by the likelihood function and the associated sensory uncertainty, both of which vary from trial to trial with the neural activity. To test this hypothesis, we decoded trial-to-trial likelihood functions from the population activity in visual cortex and used them in conjunction with a highly constrained, theoretically motivated decision model (the Bayesian model) to predict behavior. We found that a model utilizing the full likelihood function predicted the monkeys’ choices better than alternative models that ignore variations in the shape of the likelihood function. Our results provide the first population-level evidence in support of the theoretical framework of probabilistic population coding, where the same neurons that encode specific world state variables also encode the uncertainty about those variables. Importantly, under this framework the brain performs Bayesian inference under a generative model of the neural activity.

Our findings were made possible by recording from a large population simultaneously and by using a task in which uncertainty is relevant to the animal. In addition, we decoded likelihood functions using a deep neural network that does not rely on the strong parametric assumptions about the underlying generative model of the population that have dominated previous work. Importantly, we conditioned our analyses on the stimulus to rule out a confounding effect of the stimulus on the observed relationship between the decoded likelihood function and the subject’s decision. This approach is critical because previous behavioral studies on cue combination and Bayesian integration, for instance, always relied on varying stimulus features (e.g. contrast, blur, motion coherence) to manipulate uncertainty^7,8,22,38^. As a result, these studies cannot rule out that any observed correlation between a proposed method of encoding uncertainty and a subject’s behavior may be confounded by the stimulus (Supplementary Fig. 4), and they therefore fail to provide a sufficiently rigorous assessment on the representation of uncertainty. Carefully controlling for the effect of stimulus fluctuations allowed us to present rigorous evidence that the trial-by-trial fluctuations in the likelihood functions carry behaviorally relevant stimulus uncertainty information.

After showing that this likelihood function is used behaviorally, what more can we say about the neural encoding of perceptual uncertainty? First, our network learns the *log-likelihood of s*, i. e. 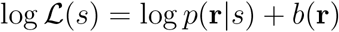 as a function of *s*. We never commit to a particular generative model *p*(**r**|*s*) as a function of **r**, as the DNN has an arbitrary offset as a function of **r** (Eq. 1 in Methods). Second, we had to move away from Poisson-like variability to better characterize the responses at the cost of analytic forms and easy interpretability. We see this as a necessary evil; namely, we have shown that making the Poisson-like assumption leads to worse predictions of behavior. That being said, the DNN extends what we know about generative models in visual cortex (e.g. tuning curves, contrast gain); in particular, it allows for rich correlation among units in the population. Third, we would like to stress that we do not believe that the DNN that we use to decode the likelihood is literally implemented in the brain. It remains an important question, and avenue for future research, what kind of transformation, if any, the brain performs in order to utilize and compute with this information.

While the sensory likelihood function is a crucial building block for probabilistic computation in the brain, fundamental questions remain regarding the nature of such computation. First, how do downstream areas process the information contained in sensory likelihood functions to make better decisions? Previous work has manually constructed neural networks for downstream computation that relied heavily on the assumption of Poisson-like variability^9,10,15,39–41^. However, more recent work has demonstrated that training generic shallow networks accomplishes the same goal without the need for task-specific manual construction^42^. Second, does each area in a feedforward chain of computation encode a likelihood function over its own variable, with the computation propagating the uncertainty information from one variable to the next? For example, in our task, it is conceivable that prefrontal cortex encodes a likelihood function over class that is derived from a likelihood function over orientation coming in from V1. Third, what are the relative contributions of feedforward, recurrent, and feedback connections to the trial-to-trial population activity and the resulting decoded likelihood functions? Some work has argued strongly for a role of feedback^28,43,44^; in the present work, we are agnostic to this issue. While answering these questions will require major efforts, we expect that our findings will help put those efforts on a more solid footing. In the meantime, our results elevate the standing of Bayesian models of perception from frameworks to describe optimal input-response mappings^45,46^ to process models whose internal building blocks—likelihood functions and probability distributions—are more concretely instantiated in neuronal activity^6,47,48^.

## Methods

### Experimental model and subject details

All behavioral and electrophysiological data were obtained from two healthy, male rhesus macaque (*Macaca mulatta*) monkeys (L and T) aged 10 and 7 years and weighting 9.5 and 15.1 kg, respectively. All experimental procedures complied with guidelines of the NIH and were approved by the Baylor College of Medicine Institutional Animal Care and Use Committee (permit number: AN-4367). Animals were housed individually in a room located adjacent to the training facility on a 12h light/dark cycle, along with around ten other monkeys permitting rich visual, olfactory, and auditory social interactions. Regular veterinary care and monitoring, balanced nutrition and environmental enrichment were provided by the Center for Comparative Medicine of Baylor College of Medicine. Surgical procedures on monkeys were conducted under general anesthesia following standard aseptic techniques.

### Stimulus presentation

Each visual stimulus was a single drifting oriented sinusoidal grating (spatial frequency: 2.79 cycles/degree visual angle, drifting speed: 3.89 cycles/s) presented through a circular aperture situated at the center of the screen. The size of the aperture was adjusted to cover receptive fields of the recorded populations, extending 2.14° and 2.86° of visual angle for Monkey L and Monkey T, respectively. The orientation and contrast of the stimulus were adjusted on a trial-by-trial basis as will be described later. The stimulus was presented on a CRT monitor (at a distance of 100 cm; resolution: 1600 × 1200 pixels; refresh rate: 100 Hz) using Psychophysics Toolbox^49^. The monitor was gamma-corrected to have a linear luminance response profile. Video cameras (DALSA genie HM640; frame rate 200Hz) with custom video eye tracking software developed in LabVIEW were used to monitor eye movements.

### Behavioral paradigm

On a given trial, the monkey viewed a drifting oriented grating with orientation θ, drawn from one of two classes, each defined by a Gaussian probability distribution. Both distributions have a mean of 0° (grating drifting horizontally rightward, positive orientation corresponding to counter-clockwise rotation), but their standard deviations differed: *σ*_1_ = 3° for class 1 (*C* =1) and *σ*_2_ = 15° for class 2 (*C* = 2). On each trial, the class was chosen randomly with equal probability, with the orientation of the stimulus then drawn from the corresponding distribution, *p*(*θ*|*C*). At the beginning of each recording session, at least three distinct values of contrasts were selected, and one of these values was chosen at random on each trial. Unlike more typical two-category tasks using distributions with identical variances but different means, optimal decision-making in our task requires the use of sensory uncertainty on a trial-by-trial basis^15^.

Each trial proceeded as follows. A trial was initiated by a beeping sound and the appearance of a fixation target (0.15° visual angle) in the center of the screen. The monkey fixated on the fixation target for 300 ms within 0.5°−1° visual angle. The stimulus then appeared at the center of the screen. After 500 ms, two colored targets (red and green) appeared to the left and the right of the grating stimulus (horizontal offset of 4.29° from the center with the target diameter of 0.71° visual angle), at which point the monkey saccaded to one of the targets to indicate their choice of class. For Monkey L, the grating stimulus was removed from the screen when the saccade target appeared, while for Monkey T, the grating stimulus remained on the screen until the subject completed the task by saccading to the target. The left-right configuration of the colored targets were varied randomly for each trial. Through training, the monkey learned to associate the red and the green targets with the narrow (*C* =1) and the wide (*C* = 2) class distributions, respectively. For illustrative clarity, we used blue to indicate *C* = 2 throughout this document. The monkey received a juice reward for each correct response (0.10−0.15 mL).

During the training, the monkeys were first trained to perform the colored version of the task, where the grating stimulus was colored to match the correct class *C* for that trial (red for *C* =1 and green for *C* = 2). Under this arrangement, the monkey simply learned to saccade to the target matching the color of the grating stimulus, although the grating stimulus orientation information was always present. As the training proceeded, we gradually removed the color from the stimulus, encouraging the monkey to make use of the orientation information in the stimulus to perform the task. Eventually, the color was completely removed, and at that point the monkey was performing the full version of the task.

### Surgical Methods

Our surgical procedures followed a previously established approach^28,50,51^. Briefly, a custom-built titanium cranial headpost was first implanted for head stabilization under general anesthesia using aseptic conditions in a dedicated operating room. After premedication with Dexamethasone (0.25-0.5 mg/kg; 48 h, 24 h and on the day of the procedure) and atropine (0.05 mg/kg prior to sedation), animals were sedated with a mixture of ketamine (10 mg/kg) and xylazine (0.5 mg/kg). During the surgery, anesthesia was maintained using isoflurane (0.5-2%). After the monkey was fully trained, we implanted a 96-electrode microelectrode array (Utah array, Blackrock Microsystems, Salt Lake City, UT, USA) with a shaft length of 1 mm over parafoveal area V1 on the right hemisphere. This surgery was performed under identical conditions as described for headpost implantation. To ameliorate pain, analgesics were given for 7 days following a surgery.

### Electrophysiological recording and data processing

The neural signals were pre-amplified at the head stage by unity gain preamplifiers (HS-27, Neuralynx, Bozeman MT, USA). These signals were then digitized by 24-bit analog data acquisition cards with 30 dB onboard gain (PXI-4498, National Instruments, Austin, TX) and sampled at 32 kHz. Broadband signals (0.5 Hz to 16 kHz) were continuously recorded using custom-built LabVIEW software for the duration of the experiment. Eye positions were tracked at 200 Hz using video cameras (DALSA genie HM640) with custom video eye tracking software developed in LabVIEW. The spike detection was performed offline according to a previously described method^26,28,50^. Briefly, a spike was detected when the signal on a given electrode crossed a threshold of five times the standard deviation of the corresponding electrode. To avoid artificial inflation of the threshold in the presence of a large number of high-amplitude spikes, we used a robust estimator of the standard deviation^52^, given by median(|*x*|)/0.6745. Spikes were aligned to the center of mass of the continuous waveform segment above half the peak amplitude. Code for spike detection is available online at https://github.com/atlab/spikedetection. In this study, the term “multiunit” refers to the set of all spikes detected from a single channel (i.e. electrode) of the Utah array, and all analyses in the main text were performed on multiunits. For each multiunit, the total number of spikes during the 500 ms of pre-target stimulus presentation, *r_i_* for the *i*^th^ unit, was used as the measure of the multiunit’s response for a single trial. The population response **r** is the vector of spike counts for all 96 multiunits.

### Dataset and inclusion criteria

We recorded a total of 61 and 71 sessions from Monkey L and T, for a total of 112,072 and 193,629 trials, respectively. We removed any trials with electrophysiology recordings contaminated by noise in the recording devices (e.g. poor grounding connector resulting in movement noise) or equipment failures. To do so, we established the following trial inclusion criteria:

1. The total spike counts 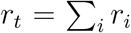 across all channels should fall within the ±4*σ*_adj_ from the median total spike counts across all trials from a single session. *σ*_adj_ is the standard deviation of the total spike count distribution robustly approximated using the interquartile range IQR as follows: 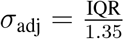.
2. For at least 50% of all units, the observed *i*^th^ unit spike count *r_i_* for the trial should fall within a range defined as: |*r_i_* − MED*_i_*| ≤ 1.5 · IQR*_i_*, where MED*_i_* and IQR*_i_* are the median and interquartile ranges of the *i*^th^ unit spike counts distribution throughout the session, respectively.

We only included trials that satisfied both of the above criteria in our analysis. Empirically, we found the above criteria to be effective in catching obvious anomalies in the spike data while introducing minimal bias into the data. After the application of the criteria, we were left with 110,695 and 192,631 trials for Monkey L and T, thus retaining 98.77% and 99.48% of the total trials, respectively. While this selection criteria allowed us to remove apparent anomaly in the data, we found that the main findings described in this paper were not sensitive to the precise definition of the inclusion criteria.

During each recording session, stimuli were presented under three or more contrast values. In all analyses to follow, we studied the trials from distinct contrast separately for each recording session, and we refer to this grouping as a “contrast-session”.

### Receptive field mapping

On the first recording session for each monkey, the receptive field was mapped using spike-triggered averaging of the multiunit responses to a white noise random dot stimulus. The white noise stimulus consisted of square dots of size 0.29° of visual angle presented on a uniform gray background, with randomly varying location and color (black or white) every 30 ms for 1 second. We adjusted the size of the grating stimulus as necessary to ensure that the stimulus covers the population receptive field entirely.

### Full-Likelihood decoder

Given the population activity **r** in response to an orientation *θ*, we aimed to decode uncertainty information in the form of a likelihood function 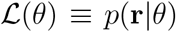, as a function of *θ*. This may be computed through the knowledge of the generative relation leading from *θ* to **r**—that is, by describing the underlying orientation conditioned probability distribution over **r**, *p*(**r**|*θ*). This procedure is typically approximated by making rather strong assumptions about the form of the density function, for example by assuming that neurons fire independently and each neuron fires according to the Poisson distribution^19^. Under this approach, the expected firing rates (i.e. tuning curves) of the *i*^th^ neuron E[*r_i_*|*θ*] = *f_i_*(*θ*) must be approximated as well, for example by fitting a parametric function (e.g. von Mises tuning curves^53^) or employing kernel regression^19^. While these approaches have proven useful, the effect of the strong and likely inaccurate assumptions on the decoded likelihood function remains unclear. Ideally, we can more directly estimate the likelihood function 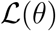 without having to make strong assumptions about the underlying conditional probability distribution over **r**.

To this end, we employed a deep neural network (DNN)^16^ to directly approximate the likelihood function over the stimulus orientation, *θ*, from the recorded population response **r**. Here we present a brief derivation that serves as the basis of the network design and training objective. Let us assume that *m* multiunits were recorded simultaneously in a single recording session, so that **r** ∈ ℝ^m^. To make the problem tractable, we bin the stimulus orientation *θ* into n distinct values, *θ*_1_ to *θ_n_* (the derivation holds in general for arbitrarily fine binning of the orientation). With this, the likelihood function can be captured by a vector L ∈ ℝ*^n^* where 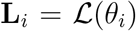. Let us assume that we can train some DNN to learn a mapping *f* from the population response **r** to the log of the likelihood function L up to a constant offset *b*. That is, *f*: ℝ*^m^* ℝ*^n^*,

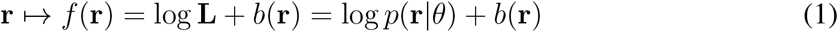

for some scalar function *b* ∈ ℝ. As the experimenter, we know the distribution of the stimulus orientation, **p**_*θ*_ ∈ ℝ*^n^*, where **p**_*θ,i*_ = **p**(*θ_i_*). We combine *f* (**r**) and **p***_θ_* to compute the log posterior over stimulus orientation *θ* up to some scalar value *b′*(**r**),

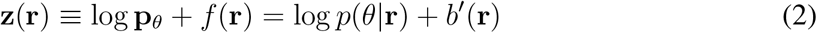

We finally take the softmax of **z**(**r**), and recover the normalized posterior function **q**(**r**) softmax(**z**(**r**)) where,

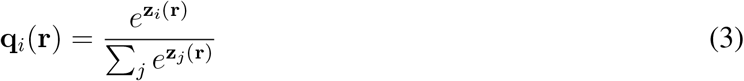

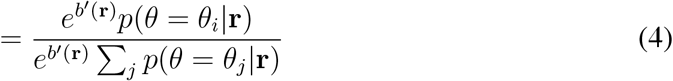

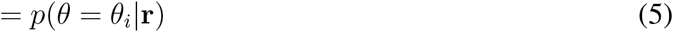

Overall, **q**(**r**) = softmax(log **p***_θ_* + *f*(**r**)).

The goal then is to train the DNN *f*(**r**) such that the overall function **q**(**r**) matches the posterior over the stimulus, **p**(**r**) where **p***_i_*(**r**) = *p*(*θ* = *θ_i_*|**r**) based on the available data. This in turn allows the network output *f*(**r**) to approach the log of the likelihood function **L**, up to a constant *b*(**r**). For 1-out-of-n classification problems, minimizing the cross-entropy between **q**(**r**) and the stimulus orientation *θ* for a given **r** lets the overall function **q**(**r**) approach the true posterior **p**(**r**), as desired^54,55^. To show this, let us start by minimizing the difference between the model estimated posterior **q**(**r**) and the true posterior **p**(**r**) over the distribution of **r**. We do this by minimizing the loss *L* defined as the expected value of the Kullback-Leibler divergence^56^ between the two posteriors:

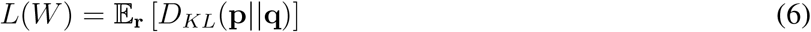

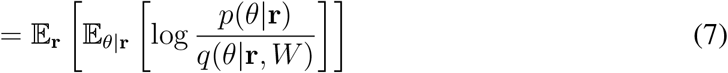

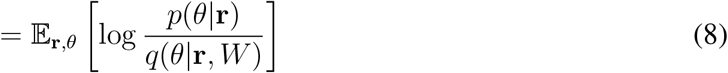

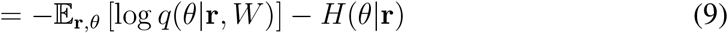

where *p*(*θ* = *θ_i_*|**r**) ≠ **p***_i_*(**r**), *q*(*θ* = *θ*_i_|**r**, *W*) ≡ **q***_i_*(**r**, *W*), *W* is a collection of all trainable parameters in the network, and *H*(*θ*|**r**) is the conditional entropy of *θ* conditioned on **r**, which is an unknown but a fixed quantity with respect to *W* and the data distribution. Here we used the notation **q**(**r**, *W*) to highlight the dependence of the network estimated posterior **q**(**r**) on the network parameters *W*. We can redefine the loss, *L*\*, only leaving the terms that depends on the trainable parameters *W*, and then apply a Monte Carlo method^57^ to approximate the loss from samples:

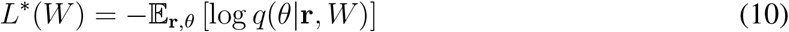

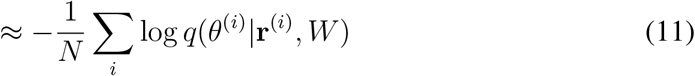

where (*θ*^(*i*)^, **r**^(*i*)^) are samples drawn from a training set for the network. Eq. 11 is precisely the definition of the cross-entropy as we set out to show.

Therefore, by optimizing the overall function **q**(**r**) to match the posterior distribution through the use of cross-entropy loss, the network output *f*(**r**) can approximate the log of the likelihood function 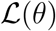 for each **r** up to an unknown constant *b*(**r**). Because we do not know the value of *b*(**r**), the network will not learn to recover the underlying generative function linking from *θ* to **r**, *p*(**r**|*θ*).

As an example, consider a neural population with responses that follows a Poisson-like distribution (i.e. a version of the exponential distribution with linear sufficient statistics^9,10^). Learning a decoder for such population responses occurs as a special case of training a DNN-based likelihood decoder. For Poisson-like variability, the stimulus-conditioned distribution over **r** is *p*(**r**|*θ*) = *ϕ*(**r**)*e*^**h**^⊤^(*θ*)**r**^. The log likelihood function is then log **L** = log *ϕ*(**r**) + **H**^⊤^**r**, where **H** is a matrix whose *i*^th^ column is **h**(*θ_i_*). If we let *f*(**r**) = **H**^⊤^**r**, then *f*(**r**) = log **L** + *b*(**r**) as desired, for *b*(**r**) = − log *ϕ*(**r**). Hence, if we used a simple fully connected network, training the network is equivalent to fitting the kernel function **h**(*θ*) of the Poisson-like distribution.

In this work, we modeled the mapping *f*(**r**) as a DNN with two hidden layers^17^, consisting of two repeating blocks of a fully connected layer of size *N_h_* followed by a rectified linear unit (ReLU)^16^ and a drop-out layer^58^ with dropout rate *d_r_*, and a fully connected readout layer with no output nonlinearity (Fig. 3c). To encourage smoother likelihood functions, we added an *L*_2_ regularizer on log **L** filtered with a Laplacian filter of the form **h** = [−0.25, 0.5, −0.25]. Therefore, the training loss included the term:

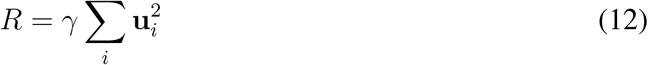

for **u** = (log **L**) * **h**, where * denotes convolution operation, **u***_i_* is the *i*^th^ element of the filtered log likelihood function **u**, and *γ* is the weight on the smoothness regularizer.

We trained a separate instance of the network for each contrast-session, and referred to this class of DNN based likelihood decoder as the Full-Likelihood decoder to differentiate from alternative decoders described later.

During the training, each contrast-session was randomly split in proportions of 80% / 20% to yield the training set and the validation set, respectively. The stimulus orientation *θ* was binned into integers in the range [−45°, 45°], and we excluded trials with orientations outside this range. This led to the exclusion of 157 out of 110,695 trials (0.14%) and 254 out of 192,631 trials (0.13%) for Monkey L and T data, respectively. The network was trained on the training set, starting with initial learning rate of λ_0_ and its performance on the validation set was monitored to perform early stopping^59^, and subsequently hyperparameter selection. For early stopping, we computed the mean squared error (MSE) between the maximum-a-posteriori (MAP) readout of the network output posterior **q** and the stimulus orientation *θ* on the validation set, and the training under a particular learning rate was terminated (early-stopped) if MSE failed to improve over 400 consecutive epochs, where each epoch is defined as one full pass through the training set. Upon early stopping, the parameter set that yielded the best validation set MSE during the course of the training was restored. The restored network was then trained again but with an updated learning rate 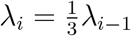, employing the same early stopping criteria. This procedure was repeated 4 times, therefore training the network under the 4 sequentially decreasing learning rate schedule of 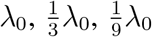 and 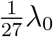. Once the training was complete, the trained network was evaluated on the validation set to yield the final score, which served as the basis for our hyperparameter selections. The values of hyperparameters for the networks, including the size of the hidden layers *N_h_*, the initial learning rate λ_0_, the weight on the likelihood function smoothness regularizer *γ*, and the drop-out rate *d_r_* during the training were selected by performing a random grid search over candidate values to find the combination that yielded the best validation set score for each contrast-session instance of the network (Supplementary Table 1). We observed that all possible values of hyperparameters were found among the best selected hyperparameter networks across all contrast-sessions and all types of networks trained.

### Decoding likelihood functions from known response distributions

To assess the effectiveness of the DNN-based likelihood decoding method described above, we simulated neural population responses with known noise distributions, trained DNN decoders on the simulated population responses, and compared the decoded likelihood functions to the ground-truth likelihood functions obtained by inverting the known generative model for the responses. We also compared the quality of the DNN-decoded likelihood functions to those decoded by assuming independent Poisson distribution on the population responses, as done in previous work^14,18,19,21,22^.

We simulated the activities of a population of 96 multiunits **r**_sim_ responding to the stimulus orientation *θ* drawn from the the distribution defined for our task such that:

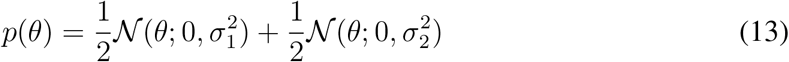

where *σ*_1_ = 3° and *σ*_2_ = 15°.

We modeled the expected response of *i*^th^ unit to *θ*—that is, the tuning function *f_i_*(*θ*)—with a Gaussian function:

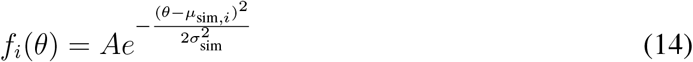

For the simulation, we have set *A* = 6 and *σ*_sim_ = 21°. We let the mean of the Gaussian tuningcurves for the 96 units to uniformly tile the stimulus orientation between −40° and 40°. In other words,

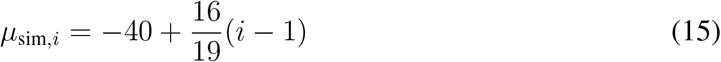

for *i* ∈ [1, 96].

For any given trial with a drawn orientation *θ*, the population response ***r***_sim_ was then generated under two distinct models of distributions. In the first case, the population responses were drawn from an independent Poisson distribution as is commonly assumed in many works:

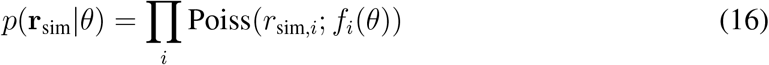

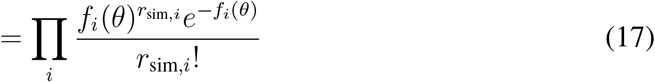

In the second case, the population responses were drawn from a multivariate Gaussian distribution with covariance matrix 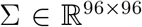 that scales with the mean response of the population.

That is:

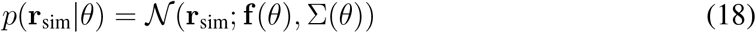

for

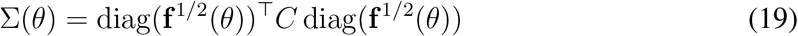

where **f**^1/2^(*θ*) ∈ ℝ^96^ such that 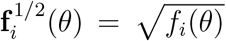, and *C* ∈ ℝ^96×96^ is a correlation matrix. Under this distribution, the variance of any unit’s response scales linearly with its mean just as in the case of the Poisson distribution, but the population responses can be highly correlated depending on the choice of the correlation matrix *C*. For the simulation, we randomly generated a correlation matrix with the average units correlation of 0.227.

For each case of the distribution, we simulated population responses for the total of 1200 trials. Among these, 200 trials were set aside as the test set. We trained the DNN-based likelihood decoder on the remaining 1000 trials, splitting them further into 800 and 200 trials as the training and validation set, respectively. We followed the exact DNN training and hyperparameter selection procedure as described earlier.

For comparison, we also decoded the likelihood function from the population response **r**_sim_ under the assumption of independent Poisson variability, regardless of the “true” distribution. Each unit’s responses over the 1000 trials were fitted separately with a Gaussian tuning curve (Eq. 14). The parameters of the tuning curve *A_i_*, *μ_i_* and *σ*_sim, i_ were obtained by minimizing the least square difference between the Gaussian tuning curve and the observed *i*^th^ unit’s responses (*θ*, *r*_sim, *i*_) using `least_squares` function from Python SciPy optimization library.

The ground-truth likelihood function *p*(**r**_sim_|*θ*) was computed for each simulated trial according to the definition of the distribution as found in Eq. 16 for the independent Poisson population or Eq. 18 for the mean scaled correlated Gaussian population.

We then assessed the quality of the decoded likelihood functions under the independent Poisson model 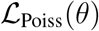 and under the DNN model **L**_DNN_ by computing their Kullback-Leibler (KL) divergence^56^ from the ground-truth likelihood function 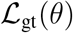, giving rise to *D*_Poiss_ and *D*_DNN_, respectively. All continuous likelihood functions (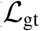 and 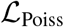) were sampled at orientation *θ* where *θ* ∈ ℤ and *θ* ∈ [−45°, 45°], giving rise to discretized likelihood functions **L**_gt_ and **L**_Poiss_ matching the dimensionality of the discretized likelihood function **L**_DNN_ computed by the DNN. We then computed the KL divergence as:

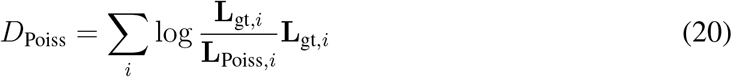

and

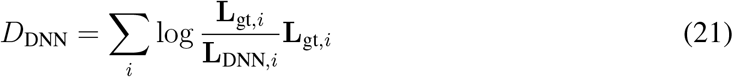

We computed the KL divergence for both models across all 200 trials in the test set for both simulated population distributions (Supplementary Fig. 3). When the simulated population distribution was independent Poisson, then *D*_Poiss_ < *D*_DNN_ for all test set trials, indicating that **L**_Poiss_ better approximated **L**_gt_ overall than **L**_DNN_. However, **L**_DNN_ still closely approximated **L**_gt_.

When the simulated population distribution was mean scaled correlated Gaussian, **L**_DNN_ better approximated **L**_gt_ than **L**_Poiss_ on the majority of the trials. Furthermore, **L**_Poiss_ provided qualitatively worse fit to the **L**_gt_ for the simulated correlated Gaussian distribution compared to the fit of **L**_DNN_ to **L**_gt_ for the simulated independent Poisson distribution.

Overall, the simulation results suggest that (1) when the form of the underlying population distribution is known (such as in the case of independent Poisson population), more accurate likelihood functions can be decoded by directly using the knowledge of the population distribution than through the DNN-based likelihood decoder, but (2) when the form of the underlying distribution is unknown (such as in the case of the mean scaled correlated Gaussian distribution), then a DNN-based likelihood decoder can yield much more accurate likelihood functions than if one was to employ a wrong assumption about the underlying distribution in decoding likelihood functions, and (3) a DNN-based likelihood decoder can provide reasonable estimate of the likelihood function across wide range of underlying distributions. Because the true underlying population distribution is hardly ever known to the experimenter, we believe that our DNN-based likelihood decoder stands as the most flexible method in decoding likelihood functions from the population responses to stimuli.

### Fixed-Uncertainty likelihood decoder

To test whether the trial-by-trial fluctuations in the shape of the likelihood function convey behaviorally relevant information, we developed the Fixed-Uncertainty likelihood decoder — a neural network based likelihood decoder that learns a fixed shape likelihood function whose location is shifted based on the input population response.

The Fixed-Uncertainty decoder network consisted of two parts: a learned fixed shape likelihood function **L**_0_ and a DNN that reads out a single scalar value Δ*_s_* corresponding to the shift that is applied to **L**_0_ (Supplementary Fig. 5) to yield the final likelihood function **L**. The DNN consisted of two repeating blocks of a fully connected layer followed by ReLU and a drop-out layer, and a final fully connected readout layer with no output nonlinearity, much like the DNN used for the Full-Likelihood decoder. The log **L**_0_ was shifted by Δ*_s_* utilizing linear interpolation based grid-sampling^60^ to shift the log-likelihood function in a manner that allows for the gradient of the loss to flow back to both the shift value Δ*s* (and therefore to the DNN parameters) as well as to the likelihood function shape **L**_0_.

The output shifted log-likelihood function was then trained in an identical manner to the full-likelihood decoder described earlier, utilizing the same set of training paradigm with early stopping and regularizers, and explored the same range of hyperparameters.

### Likelihood functions based on Poisson-like and independent Poisson distributions

To serve as a comparison, for each contrast-session, we decoded likelihood functions from the population response assuming Poisson-like or independent Poisson distribution for *p*(**r**|*θ*) (Supplementary Fig. 2).

As was noted above, decoding likelihood function under the Poisson-like distribution is a special case of the Full-Likelihood decoder but using entirely linear DNN (i.e. no nonlinearity utilized in the network). Therefore, to decode likelihood functions under the assumption of the Poisson-like distribution, for each contrast-session, we trained a DNN with two hidden layers consisting of two repeating blocks of a fully connected layer followed by a drop-out layer^58^ but with no nonlinear activation functions, and a fully connected readout layer with no output nonlinearity. The rest of the training and model selection procedure was identical to that of the Full-Likelihood or the Fixed-Uncertainty decoder described earlier.

To decode likelihood function under the independent Poisson distribution assumption, we first fitted tuning curves *f_i_*(*θ*) for each multiunit’s responses to stimulus orientations *θ* within a single contrast session. Tuning curves were computed using Gaussian process regression^61^ with squared-exponential covariance function 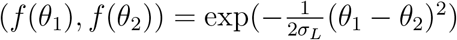 and a fixed observational noise *σ_o_* using values of *σ_L_* = 20 and *σ_o_* = 2 selected based on the cross validation performance on multiunit’s response prediction on a dataset not included elsewhere in the analysis. Once tuning curves were computed, the likelihood function over stimulus orientations was computed from the population response **r** as follows:

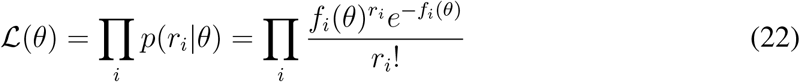

### Mean and standard deviation of likelihood function

For uses in the subsequent analyses, we computed the mean and the standard deviation of the likelihood function by treating the likelihood function as an unnormalized probability distribution:

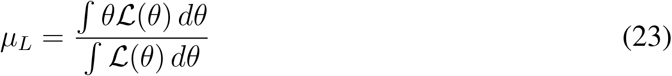

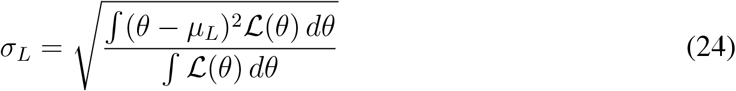

We took the *μ_L_* and *σ_L_* to be the point estimate of the stimulus orientation and the measure of the spread of the likelihood function, respectively, used in all subsequent analyses. Although not presented here, we performed the following analyses with other point estimates of the stimulus orientation such as the orientation at the maximum of the likelihood function and the median of the likelihood functions, and observed that models with mean of the likelihood function as the point estimate performed the best.

### Attribution analysis

To assess whether the same members of the population simulatenously encode the best point estimate (i.e. in the form of the mean of the likelihood function *μ_L_*) and uncertainty (i.e. in the form of the width of the likelihood function *σ_L_*), we computed the attribution of each multiunit input of the trained Full-Likelihood decoder to the mean of the likelihood *μ_L_* and the standard deviation of the likelihood function *σ_L_* giving rise to the attribution *A_μ_*, *A_σ_* ∈ ℝ^*m*^, respectively, where *m* is the number of multiunits in the input to the network. Among numerous methods of computing attribution^33–35,62^, we have selected three popular gradient based attribution methods^33^: saliency maps^34^, gradient × input^62^, and DeepLift^35^ and compared their results.

Given a collection of input population responses and computed likelihood functions {**r**^(*i*)^, **L**^(*i*)^}, where the superscript denotes the *i*^th^ trial in the contrast session, we compute the mean and the standard deviation of the likelihood function according to Eq. 23 and Eq. 24, respectively, giving rise to 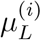 and 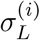. Given a target feature *S* ∈ {*μ_L_*, *σ_L_*} that can be computed from the input units **r** through a differentiable function, we compute the attribution of the input units to the target *S* for each trial according to each attribution method, yielding 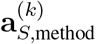, where **a** ∈ **R**^*m*^. The sign of the attribution indicates whether increasing the unit tends to increase or decrease the target feature. Since we are interested more in how much each unit contribute to the target feature rather than in which direction, we take the absolute value of per trial attribution and compute the average across all trials to yield the final attribution of the input units:

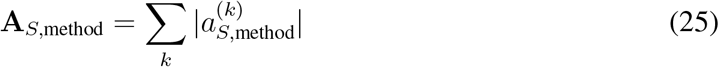

For the saliency maps based method^34^, the attribution is computed as the partial derivative of the feature *S* with respect to the input units **r**:

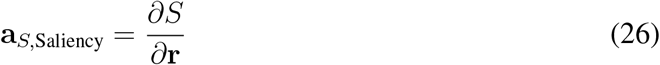

which can be computed rather straightforwardly on a DNN implemented using any of the modern neural network libraries equipped with automatic gradient computation.

For Gradient × Input (GI) method, the attribution is computed as the gradient of the feature with respect to the input (as in saliency maps) multiplied with the input **r**:

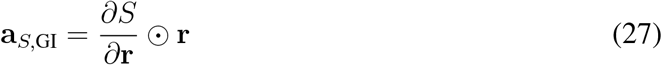

Finally, we computed DeepLIFT attribution by using modified gradient computation for ReLU units in the network defined as:

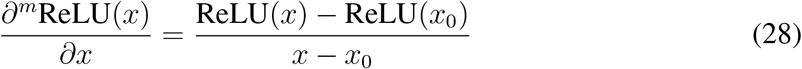

where *x*_0_ represents the input into the ReLU nonlinearity when a reference input **r**_0_ was used as the input into the network. Here, we have defined the reference network input to be the average population response across all trials (refer to Ref^33,35^ for details).

Using the above modified gradient computation for ReLU nonlinearity in the backpropagation to compute the partial derivative of the target feature with respect to the input units yield the modified partial derivative 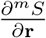 which is finally used to compute the DeepLIFT (DL) attribution as:

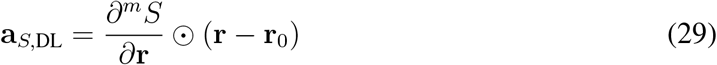

For each contrast session and each attribution method, we computed the attribution of the input units to both *μ_L_* and *σ_L_*, yielding vectors **A**_*μ*_ and **A**_*σ*_, and we computed the Pearson correlation coefficient between the two scores across the units (Fig. 6).

### Decision-making models

Given the hypothesized representation of the stimulus and its uncertainty in the form of the likelihood function 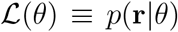, the monkey’s trial-by-trial decisions were modeled based on the assumption that the monkey computes the posterior probability over the two classes *C* = 1 and *C* = 2, and utilizes this information in making decisions—that is, in accordance to a model of a Bayesian decision maker. The orientation distributions for the two classes are 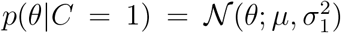 and 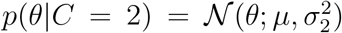 with *μ* = 0 and *σ*_1_ = 3° and *σ*_2_ = 15° where 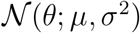 denotes a Gaussian distribution over *θ* with mean *μ* and variance *σ*^2^. The posterior ratio *ρ* for the two classes is:

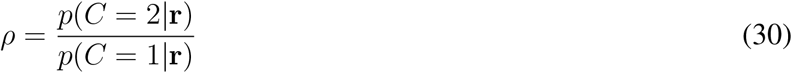

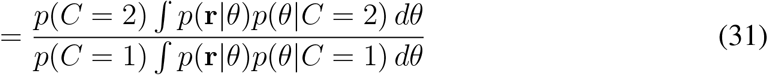

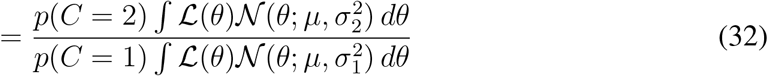

A Bayes-optimal observer should select the class with the higher probability—a strategy known as maximum-a-posteriori (MAP) decision-making:

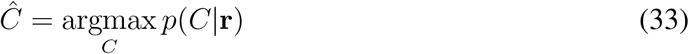

where 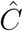 is the subject’s decision. However, according to the posterior probability matching strategy^63,64^, the decision of subjects on certain tasks are better modeled as sampling from the posterior probability:

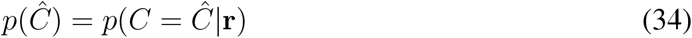

To capture either decision-making strategy, we modeled the subject’s classification decision probability ratio as follows:

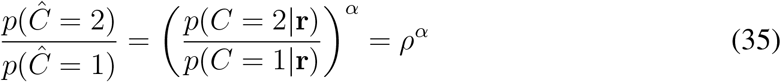

where *α* ∈ ℝ^+^. When *α* = 1, the decision-making strategy corresponds to the posterior probability matching while *α* = ∞ corresponds to the MAP strategy^64^. We fitted the value of *α* for each contrast-session during the model fitting to capture any variation of the strategy. Furthermore, we incorporated a lapse rate λ, a fraction of trials on which the subject does not pay attention and makes a random decision. Hence, the final probability that the subject selects the class *C* = 1 was modeled as:

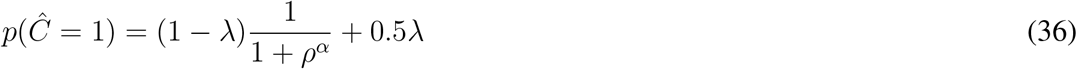

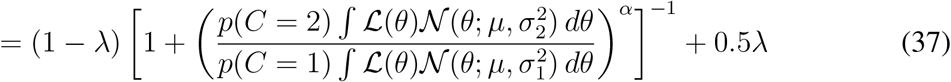

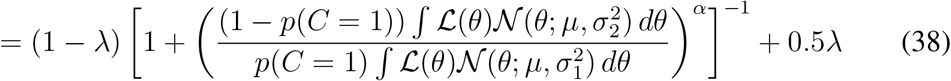

For each contrast-session, we fitted the above Bayesian decision model to the monkey’s decisions by fitting the four parameters: *μ*, *p*(*C* = 1), *α*, and λ. Fitting *μ* (the center of stimulus orientation distributions) and *p*(*C* = 1) (prior over class) allowed us to capture the bias in the stimulation distribution that the subject may have acquired errorneously during the training, and fitting *α* and λ allowed for the model to match the decision-making strategy employed by the subject.

Utilizing the likelihood function 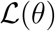 decoded from the V1 population response via the Full-Likelihood decoder network in Eq. 38 gave rise to the Full-Likelihood Model that made use of all information including the trial-by-trial uncertainty information as captured by the trial-by-trial fluctuations in the shape of the likelihood function. Alternatively, utilizing the likelihood function decoded by the trained Fixed-Uncertainty decoder gave rise to the Fixed-Uncertainty Model. The Fixed-Uncertainty Model effectively ignores all trial-by-trial fluctuations in the uncertainty that would be captured by the flucutations in the shape of the likelihood function, but captures the trial-by-trial point estimate of the stimulus orientation 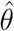 by shifting the leaned fixed shape likelihood function over orientation. For each contrast-session, different fixed likelihood shape was learned, allowing the overt measure of uncertainty such as contrast to modulate the expected level of uncertainty.

For comparison, we have also tested the performance of the trial-by-trial decision prediction utilizing likelihood functions decoded based on Poisson-like or independent Poisson population distribution assumptions, giving rise to the Poisson-like Model and the Independent Poisson Model for predicting trial-by-trial decisions, respectively.

### Model fitting and model comparison

We used 10-fold cross-validation to fit and evaluate both decision models, separately for each contrast-session. We divided all trials from a given contrast-session randomly into 10 equally sized subsets, *B*_1_, *B*_2_,…, *B*_i_,…, *B*_10_ where *B*_i_ is the *i*^th^ subset. We then held out a single subset *B_i_* as the test set, and trained the decision-making model on the remaining 9 subsets combined together, serving as the training set. The predictions and the performance of the trained model on the held out test set B_i_ was then reported. We repeated this 10 times, iterating through each subset as the test set, training on the remaining subsets.

The decision models were trained to minimize the negative log likelihood on the subject’s decision across all trials in the training set:

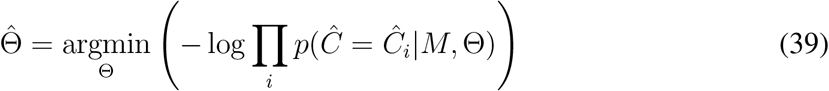

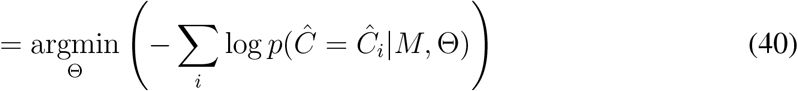

where Θ is the collection of the parameters for the decision-making model *M* and 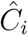 is the subject’s decision on the *i*^th^ trial in the training set. The term 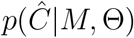 is given by the Eq. 38 with either the unmodified 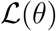 in the Full-Likelihood Model or a Gaussian approximation to 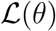 in the Fixed-Uncertainty Model.

The optimizations were performed using three algorithms: `fmincon` and `ga` from MAT-LAB’s optimization toolbox and Bayesian Adaptive Direct Search (BADS)^65^. When applicable, the optimization was repeated with 50 or more random parameter initializations. For each crossvalidation fold, we retained the parameter combination 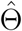 that yielded the best training set score (i.e. lowest negative log likelihood) among all optimization runs across different algorithms and parameter initializations. We subsequently tested the model *M* with the best training set parameter 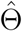 and reported the score on the test set. For each contrast-session, all analyses on the trained model presented in the main text were performed on the aggregated test sets scores.

### Likelihood shuffling analysis

To assess the contribution of the trial-by-trial fluctuations in the decoded likelihood functions in predicting the animal’s decisions under the Full-Likelihood Model, for each contrast-session we shuffled the likelihood functions among trials in the same stimulus orientation bin, while maintaining the trial to trial relationship between the point estimate of the stimulus orientation (i.e. mean of the normalized likelihood) and the perceptual decision. Specifically, we binned trials to the nearest orientation degree such that each bin was centered at an integer degree (i.e. bin center ∈ ℤ) with the bin width of 1°. We then shuffled the likelihood functions among trials in the same orientation bin. This effectively removed the stimulus orientation conditioned correlation between the likelihood function and the subject’s classification 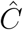, while preserving the expected likelihood function for each stimulus orientation.

However, we were specifically interested in decoupling the uncertainty information contained in the shape of the likelihood function from the decision while minimally disrupting the trial-by-trial correlation between the point estimate of the stimulus orientation and the subject’s classification decision. To achieve this, for each trial, the newly assigned likelihood function was shifted such that the mean of the normalized likelihood function, *μ_L_* (Eq. 23), remained the same for each trial before and after the likelihood shuffling (Fig. 5c). This allowed us to specifically assess the effect of distorting the shape of the likelihood function conditioned on both the (binned) stimulus orientation and the point estimate of the stimulus orientation (i.e. μL) (Fig. 5c). To ensure that both models can take the full advantage of any information that remains in the shuffled likelihood functions, we trained both the Full-Likelihood Model and the Fixed-Uncertainty Model from scratch on the shuffled data. Aside from the difference in the dataset, we followed the exact procedure used when training on the original (unshuffled) data, evaluating the performance through cross-validation on the test sets.

### Classification simulation

We computed the expected effect size of the model fit difference between the Full-Likelihood Model and the Fixed-Uncertainty Model by generating simulated data using the trained Full-Likelihood Model as the ground truth. Specifically, for each trial for each contrast-session, we computed the probability of responding 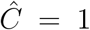 from Eq. 38, utilizing the full decoded likelihood function 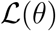 for the given trial, and sampled a classification decision from that probability. This procedure yielded simulated data where all monkeys’ decisions were replaced by decisions made by the trained Full-Likelihood Models. We repeated this procedure 5 times, thereby producing 5 sets of simulated data. For each set of simulated data, we trained the two decision-making models (Full-Likelihood Model and Fixed-Uncertainty Model) on each contrast-session with 10-fold cross-validation, and reported the aggregated test set scores as was done for the original data.

### Code availability

Code used for modeling and training the deep neural networks as well as for figure generation will be made available for view and download at https://github.com/eywalker/v1_likelihood. All other code used for analysis including data selection and decision model fitting will be placed at https://github.com/eywalker/v1_project. Finally, code used for elecrophysiology data processing can already be found in the Tolias lab GitHub organization https://github.com/atlab.

### Data availability

All figures except for Figure 1 and Supplementary Figure 4 were generated from raw data or processed data. The data generated and/or analyzed during the current study are available from the corresponding author upon reasonable request. No publicly available data was used in this study.

### Statistics

All statistical tests used were two-tailed paired two-sample t-test, unless specified otherwise. Wherever reported, data are means and error bars indicate standard error of the means computed as 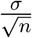 where *σ* is the standard deviation and *n* is the size of the sample within the bin, unless specified otherwise. Exact p values less than 0.001 were reported as p <0.001. When appropriate, p values were corrected for multiple comparisons and the corrected p value was reported.

## Acknowledgments

The research was supported by National Science Foundation Grant IIS-1132009 (to W.J.M. and A.S.T.), DP1 EY023176 Pioneer Grant (to A.S.T.), F30 EY025510 (to. E.Y.W.) and R01 EY026927 (to A.S.T and W.J.M.). We thank Fabian Sinz for helpful discussion and suggestions on the deep neural network fitting to likelihood functions. We also thank Tori Shin for assistance in monkeys behavioral training and experimental data collection.

## Author contributions

All authors designed the experiments and developed the theoretical framework. R.J.C. trained the first monkey, and R.J.C. and E.Y.W. recorded data from this monkey. E.Y.W. trained and recorded from the second monkey. E.Y.W. performed all data analyses. E.Y.W. wrote the manuscript, with contributions from all authors.

---

**Supplementary Figure 1:**
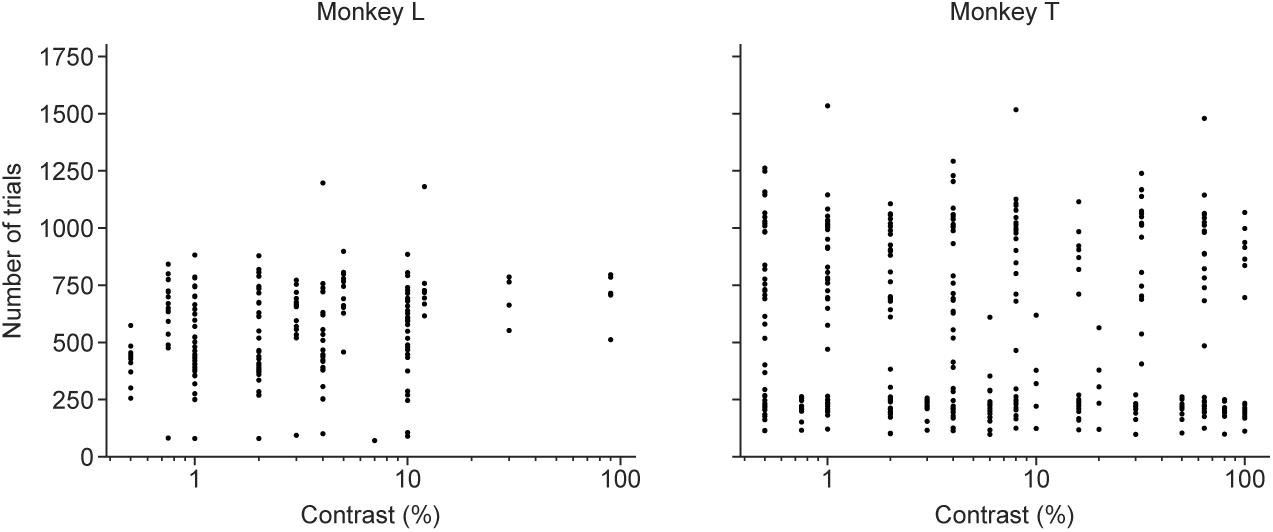
Number of trials per contrast-session. Each point corresponds to a single contrast-session, depicting the number of trials performed at the particular contrast.

**Supplementary Figure 2:**
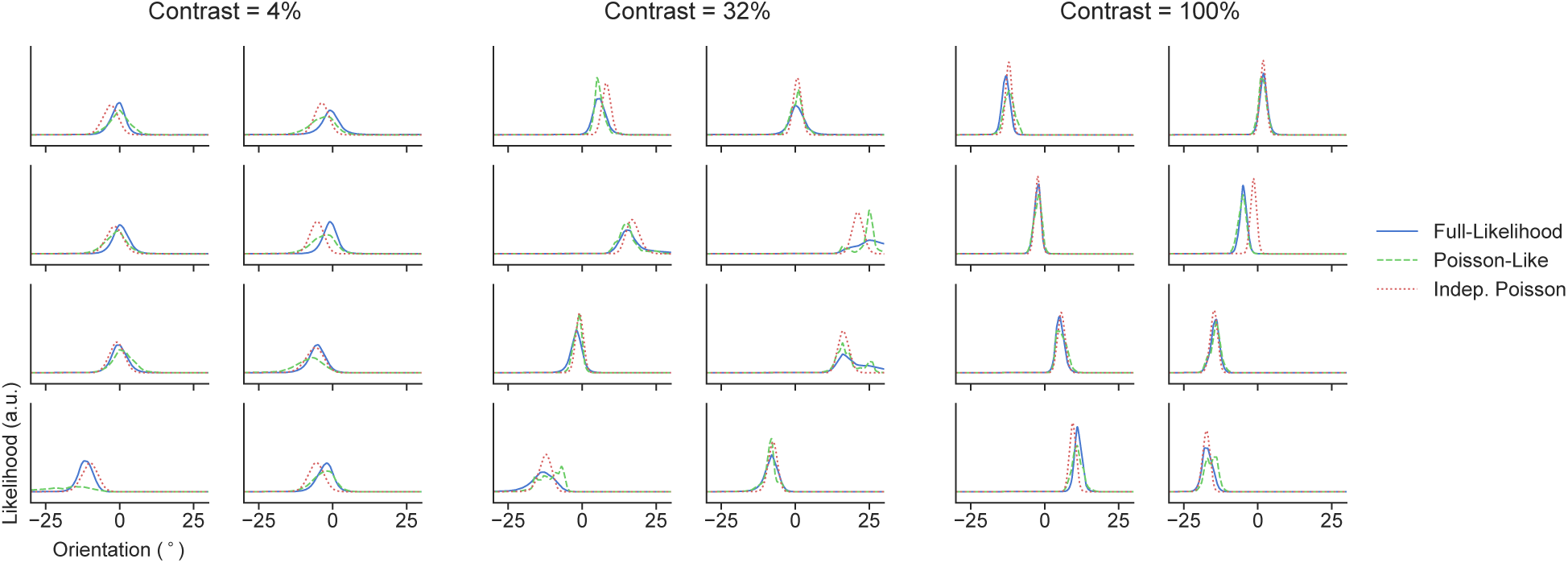
Example decoded likelihood functions. Example decoded likelihood functions under Full-Likelihood, Poisson-like and Independent-Poisson based decoders are shown for randomly selected trials from three distinct contrast-sessions from Monkey T.

**Supplementary Figure 3:**
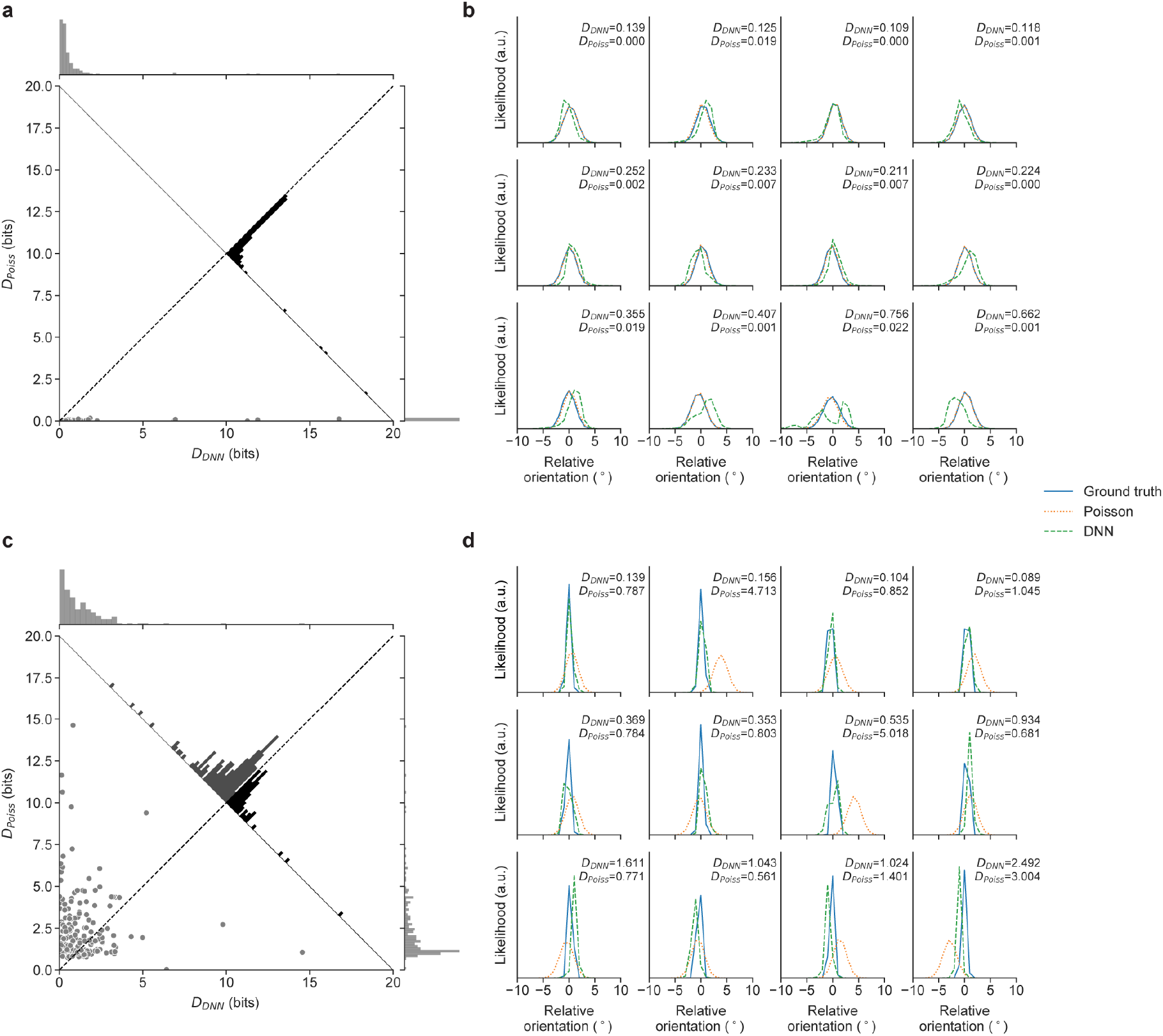
Performance of the likelihood functions decoded by DNN-based decoders. **a-b**, Results on independent Poisson population responses. **a**, KL divergence between the ground truth likelihood function and likelihood function decoded with: a trained DNN *D*_DNN_ vs. independent Poisson distribution assumption *D*_Poiss_. Each point is a single trial in the test set. The distributions of *D*_DNN_ and *D*_Poiss_ are shown at the top and right margins, respectively. The distribution of pair-wise difference between *D*_DNN_ and *D*_Poiss_ is shown on the diagonal. **b**, Example likelihood functions. The ground truth (solid blue), independent-Poisson based (dotted orange), and DNN-based (dashed green) likelihood functions are shown for selected trials from the test set. Four random samples (columns) were drawn from the top, middle and bottom 1/3 of trials sorted by the *D*_DNN_ (rows). **c-d**, Same as in **a-b** but for simulated population responses with correlated Gaussian distribution where variance is scaled by the mean.

**Supplementary Figure 4:**
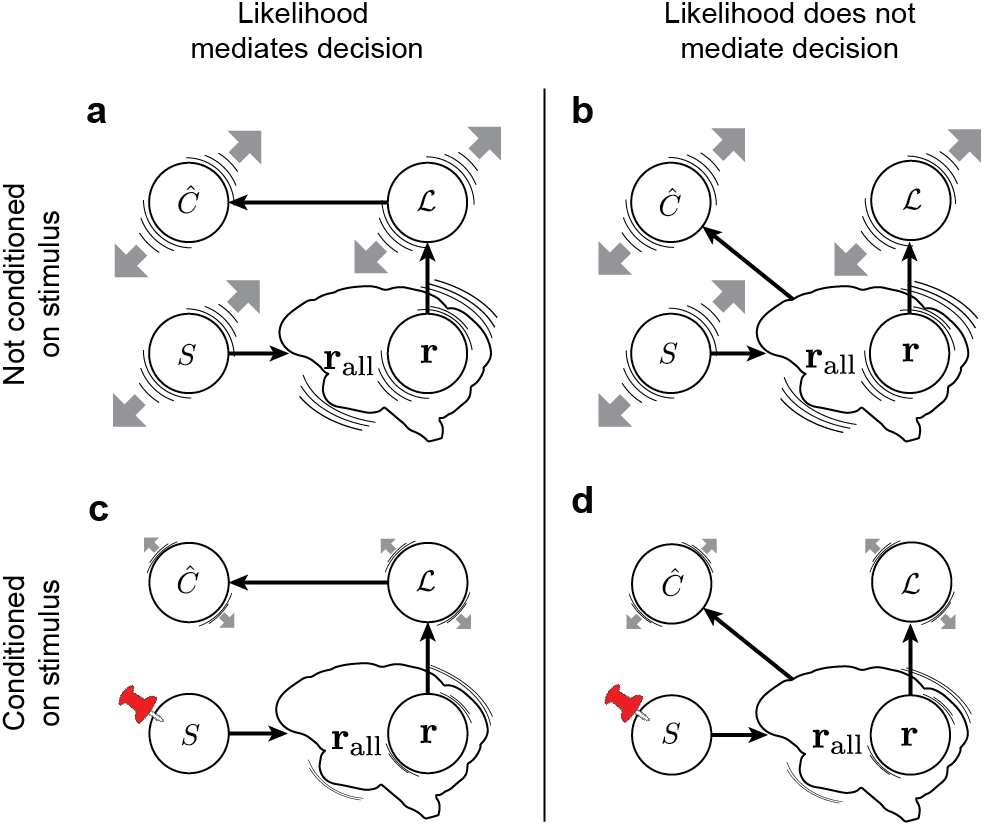
Alternative relationships between the likelihood function and the decision. Possible relationships between variables in the model are indicated by black arrows. We consider two scenarios: **a, c** the likelihood function 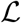 mediates the decision 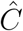, **b, d** the likelihood function does not mediate the decision. The gray arrow represents the trial-by-trial fluctuations in the subject’s decisions 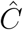 as predicted by the variable. **a, b**, When not conditioning on the stimulus *s*, the stimulus can drive correlation among all variables, making it difficult to distinguish the two scenarios. **c, d**, When conditioning on the stimulus, we expect correlation between 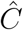 and 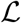 only when 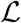 mediates the decision, allowing us to distinguish the two scenarios. The variable **r** represents the recorded cortical population and **r**_all_ represents responses of all recorded and unrecorded neurons.

**Supplementary Figure 5:**
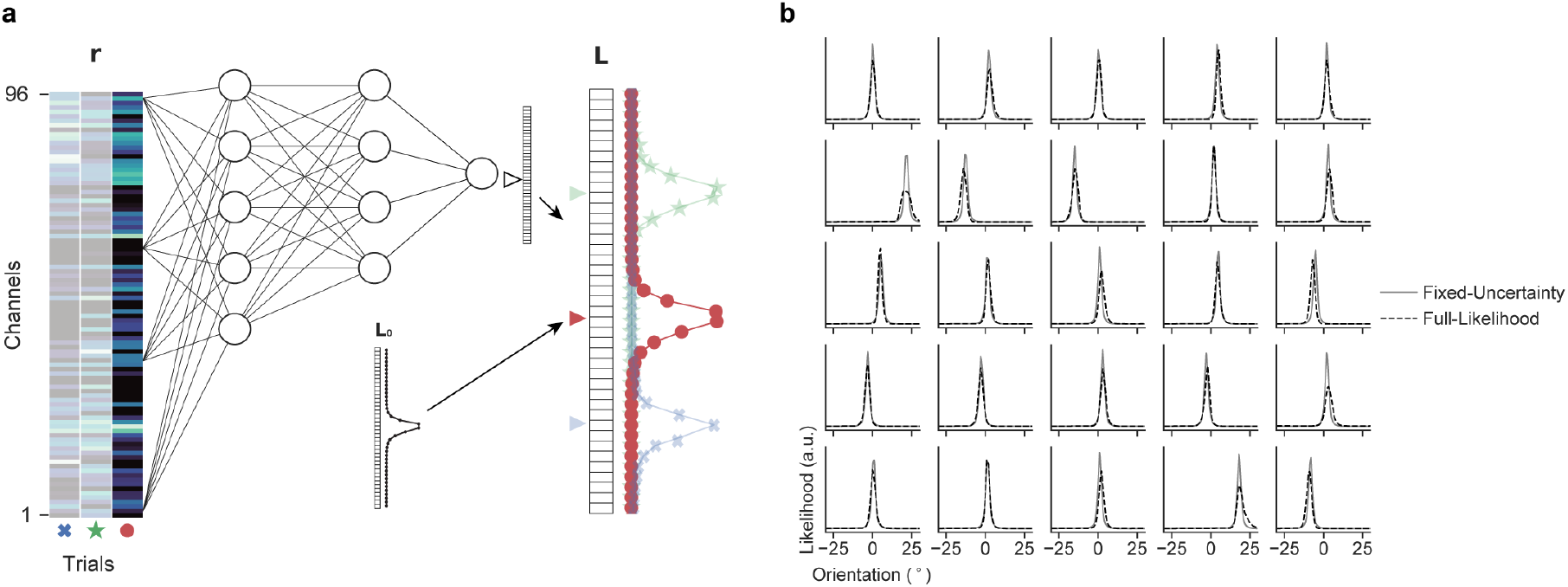
Fixed-Uncertainty decoder. **a**, A schematic of a DNN for the Fixed-Uncertainty decoder mapping **r** to the decoded likelihood function **L**. For each contrast-session, the Fixed-Uncertainty decoder learns a single fixed-shape likelihood function **L**_0_ and a network that shifts **L**_0_ based on the population response. Therefore, all resulting likelihood functions share the same shape (uncertainty) but differ in the center location from trial to trail. **b**, Example decoded likelihood functions from randomly selected trials from a single contrast session for both the Fixed-Uncertainty decoder and Full-Likelihood decoder.

**Supplementary Figure 6:**
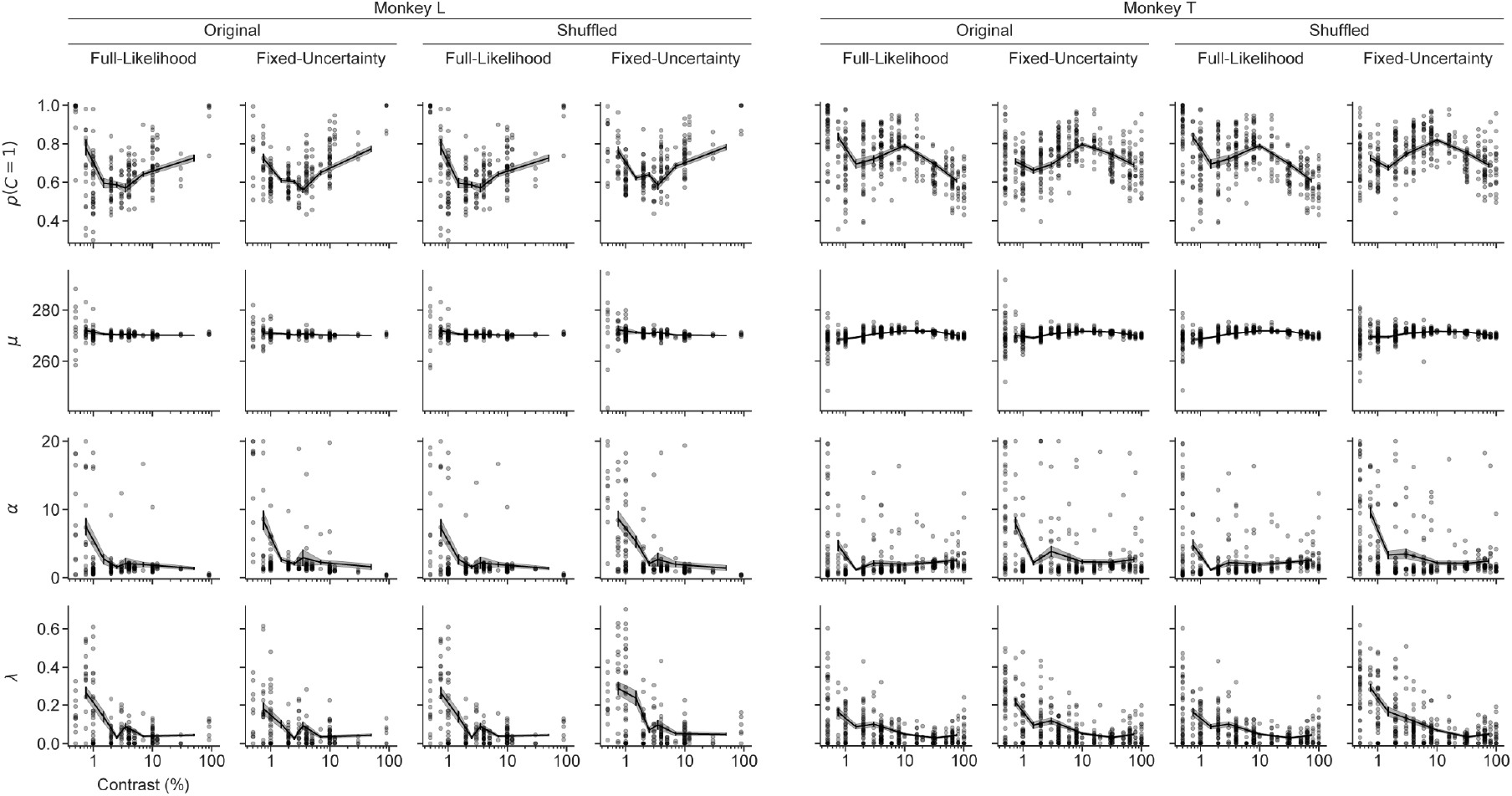
Fitted Bayesian decision model parameters. Each point corresponds to a single contrast-session, depicting the average fitted parameter value across 10 cross-validation training sets plotted against the contrast of the contrast-session. The solid line and error bars/shaded area depicts the mean and the standard error of the mean of the parameter value for binned contrast values, respectively.

**Supplementary Figure 7:**
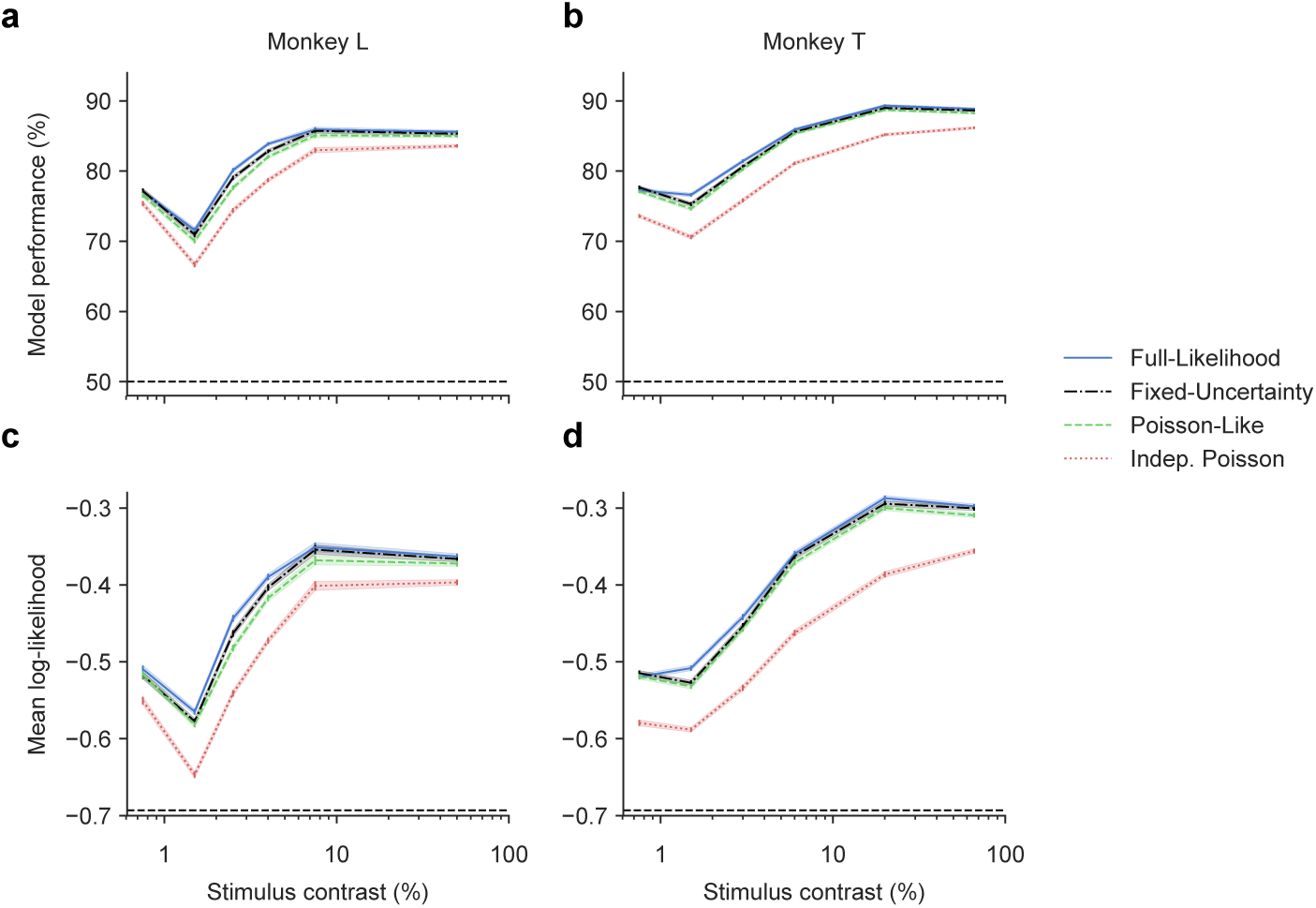
Model performance on decision predictions. **a-b**, Model performance measured in proportions of trials correctly predicted by the model as a function of contrast for four decision models based on different likelihood decoders. On each trial, the class decision that would maximize the posterior 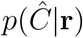 was chosen to yield a concrete classification prediction. **c-d**, Same as in **a-b** but with performance measured as the trial-averaged log likelihood of the model. For **a-b** and **c-d**, dashed lines indicate the performance at chance (50% and ln(0.5), respectively).

**Supplementary Figure 8:**
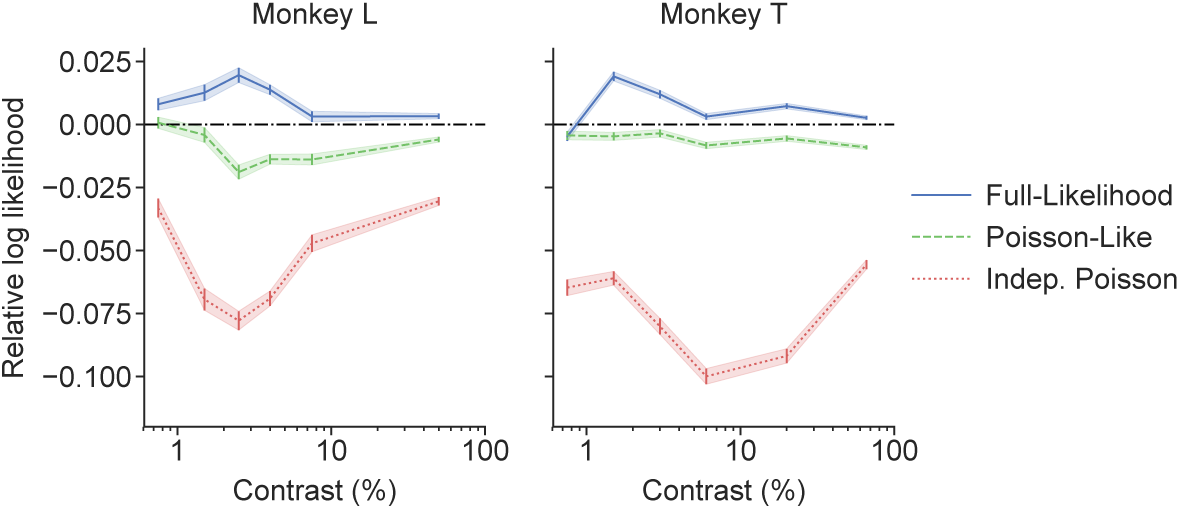
Performance of Poisson-like and Independent Poisson Models. For each monkey, the average trial-by-trial performance of the Full-Likelihood, Poisson-like and Independent Poisson Models are shown relative to the Fixed-Uncertainty Model across contrasts, measured as the average trial difference in the log likelihood.

**Supplementary Figure 9:**
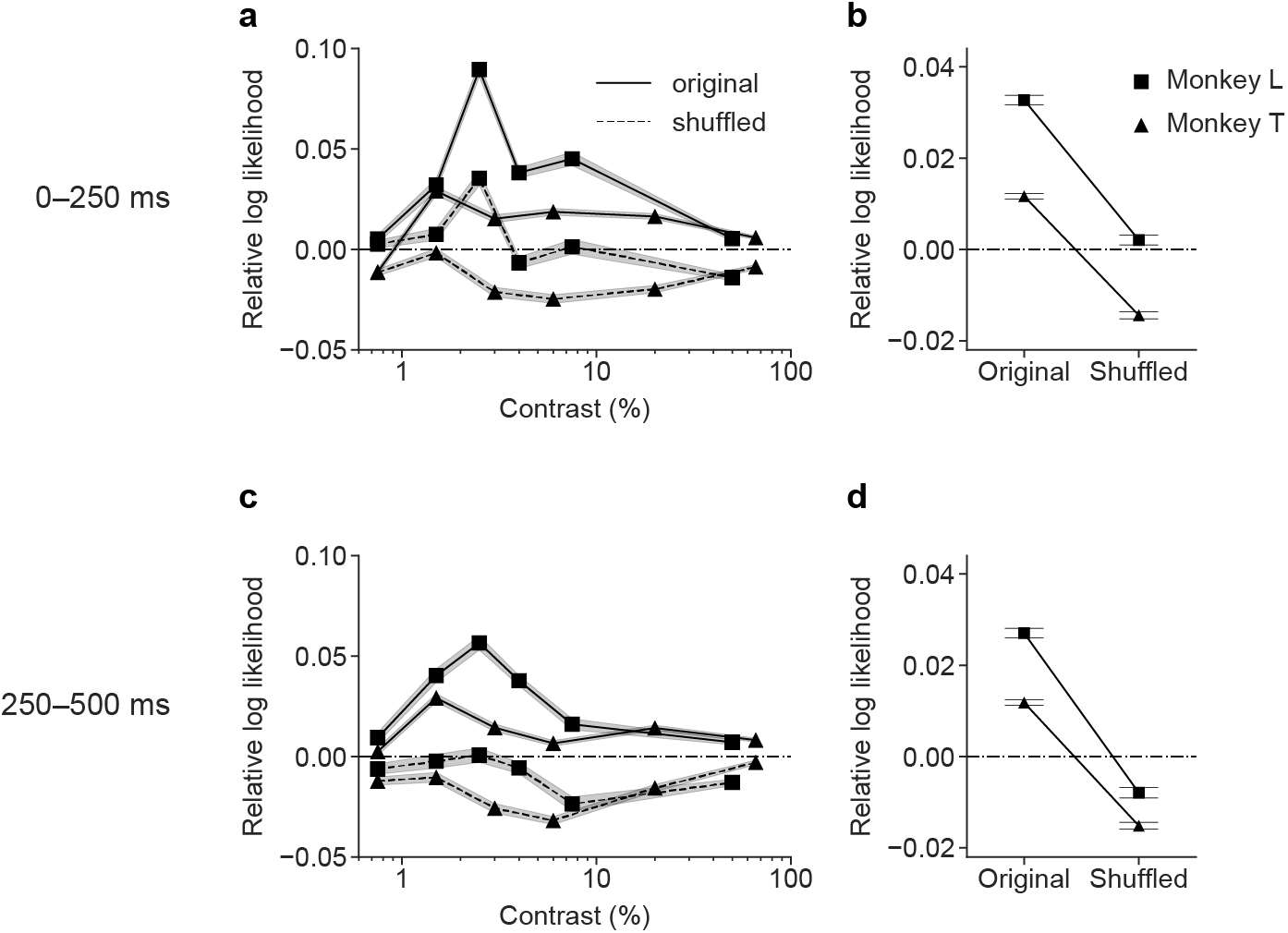
Model performance based on population responses to different stimulus windows. **a, c**, Average trial-by-trial performance of the Full-Likelihood Model relative to the Fixed-Uncertainty Model across contrasts, measured as the average trial difference in the log likelihood. The models were trained and evaluated on the population response to (**a**) the first half (0–250 ms) or (**c**) the second half (250–500 ms) of the stimulus presentation. The results for the original (unshuffled) and the shuffled data are shown in solid and dashed lines, respectively. The squares and triangles mark Monkey L and T, respectively. **b, d**, Relative model performance summarized across all contrasts based on models trained as described in (**a, c**). Performance on the original and the shuffled data is shown individually for both monkeys. The difference between the Full-Likelihood and Fixed-Uncertainty Models was significant with *p* < 0.001 for both stimulus windows, and on both the original and the shuffled data for both monkeys, except for the shuffled dataset on 0–250ms for Monkey L, for which there was no significant difference between the two models (*p* = 0.17). The difference between the Full-Likelihood Model on the original and the shuffled data was significant (*p* < 0.001 for both monkeys for both stimulus windows). For **a-d**, all data points are means, and error bar/shaded area indicate standard error of the means.

**Supplementary Figure 10:**
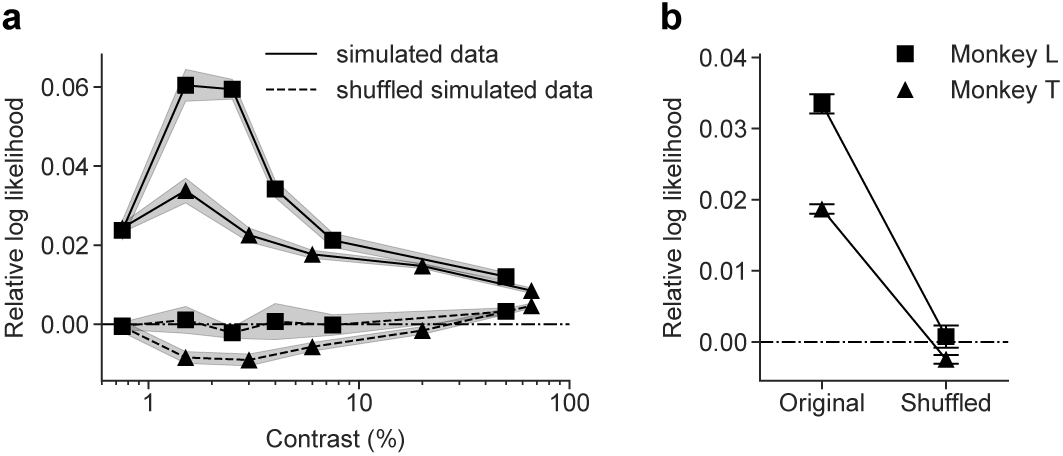
Expected model performance on simulated data using the trained Full-Likelihood Model as the ground truth. **a**, Average trial-by-trial performance of the Full-Likelihood Model relative to the Fixed-Uncertainty Model across contrasts on the simulated data, measured as the trial-averaged difference in the log likelihood. The results for the unshuffled and the shuffled simulated data are shown in solid and dashed lines, respectively. The squares and triangles mark Monkey L and T, respectively. **b**, Relative model performance summarized across all contrasts. Results are shown for each monkey and for unshuffled and shuffled simulated data. For **a** and **b**, all data points are the means and error bar/shaded area indicate the standard deviation across the 5 simulation repetitions.

**Supplementary Figure 11:**
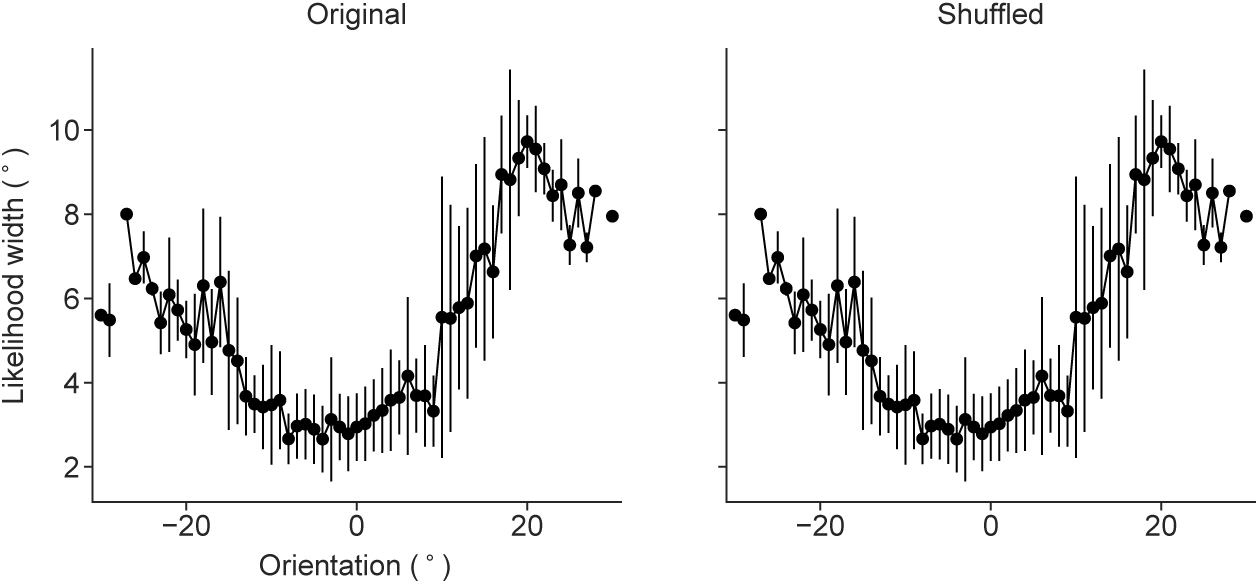
Dependence of average likelihood width on the stimulus orientation. The dependence of the width of the likelihood function *σ_L_* on the stimulus orientation is depicted for an example contrast-session (Monkey T, 8% contrast) on the original and the shuffled data. The shuffling procedure preserves the relationship between the average likelihood width and the stimulus orientation as desired.

**Supplementary Table 1:** Possible values of hyperparameters during model selection.

| Symbol | Description | Possible Values |
| --- | --- | --- |
| $N_h$ | number of hidden units per layer | $\{400, 600, 800, 1000\}$ |
| $\lambda_0$ | initial learning rate | $\{0.01, 0.03, 0.6\}$ |
| $\gamma$ | Laplacian L2 regularizer weight | $\{3, 30, 300\}$ |
| $d_r$ | dropout rate | $\{0.2, 0.5, 0.9\}$ |

